# Early tuberculosis treatment remodels blood mRNA and small RNA, and a baseline host signature is associated with early culture conversion

**DOI:** 10.64898/2026.10.25.761687

**Authors:** G V Anoushka Chinmayi, Darla Quijada, Artur T L Queiroz, Neil Martinson, Bruno B. Andrade, Marc K. Halushka, Joel S. Bader, Petros C. Karakousis

## Abstract

**Background:** Tuberculosis (TB) treatment monitoring relies on sputum-based microbiological assays, which are slow and provide limited insight into the host response. Blood-based biomarkers could offer rapid and minimally invasive treatment monitoring.

**Methods:** We performed an integrative multi-omics analysis of paired whole-blood samples collected at baseline and four weeks after treatment initiation across three South African cohorts. A meta-analysis combined RNA sequencing data from GSE89403 (86 pairs) and a prospective cohort (11 pairs), with directional concordance assessed in an independent microarray cohort, GSE31348 (27 pairs). In the prospective cohort, small RNA sequencing profiled microRNAs (miRNAs) and tRNA-derived fragments (tRFs). In GSE89403 (n = 76), a k-top scoring pairs signature (EARLY5) was developed using leave-one-out cross-validation to predict week-4 sputum culture conversion from baseline samples.

**Findings:** The meta-analysis identified 63 concordantly regulated genes. These changes reflected attenuation of neutrophil, inflammatory and complement programs, and deconvolution showed decreased neutrophils with increased T-cell and NK-cell populations. Seven miRNAs and eight nuclear-encoded tRFs changed during treatment, including erythroid-associated miR-144, with computationally identified candidate interactions involving complement genes. EARLY5 identified participants who converted by week 4 with a cross-validated AUC of 0.76 and remained associated with conversion after adjustment for baseline bacterial burden in logistic regression and performed comparably to existing TB signatures.

**Interpretation:** Early TB treatment induces coordinated remodeling of blood mRNA and small RNA profiles. Baseline host transcriptional profiles were associated with early culture conversion after accounting for bacterial burden, supporting further evaluation in independent cohorts.

**Funding:** CRDF Global/CFAR/RePORT and the National Institutes of Health (P30AI168436).

## Introduction

Tuberculosis (TB) remains the leading cause of death from a single infectious agent worldwide despite the availability of effective antimicrobial therapy ^1^. Although treatment outcomes for drug-susceptible pulmonary TB are generally favorable, response monitoring still relies largely on sputum-based microbiological methods, including smear microscopy and mycobacterial culture^1^. These methods have important limitations. Smear microscopy lacks sensitivity and cannot distinguish viable from non-viable bacilli, and culture-based methods require prolonged incubation times that delay recognition of patients with slower treatment response^1,2^. Molecular assays such as Xpert MTB/RIF may remain positive after microbiological clearance because nucleic acids from non-viable organisms persist during therapy^3^. Consequently, there is substantial interest in host-derived biomarkers that could provide rapid and minimally invasive measures of treatment response.

Whole-blood transcriptomics has emerged as a promising approach for characterizing systemic immune responses during TB infection and treatment. Active disease is associated with a neutrophil-driven interferon-inducible transcriptional signature that contracts during successful therapy, reflecting remodeling of the host response as bacterial burden declines^4^. Subsequent studies have identified increasingly parsimonious blood RNA signatures capable of distinguishing active disease from latent infection, predicting progression to active TB, and monitoring treatment response^5–9^. Transcriptional changes during treatment have also been associated with long-term clinical outcomes, suggesting that host-response dynamics may provide information beyond conventional microbiological measures^5,10^. However, most of these signatures were developed for diagnosis or risk of progression, and whether baseline host transcriptional state predicts the pace of early bacteriological response remains unclear. In addition, most studies have focused on mRNA, and the contribution of other regulatory RNA layers to treatment-associated immune remodeling is yet to be understood.

MicroRNAs (miRNAs) and tRNA-derived fragments (tRFs) are small non-coding RNAs with important roles in post-transcriptional regulation. Several miRNAs have been implicated in TB pathogenesis^11,12^, and proposed as biomarkers of disease activity and treatment response^13–17^. More recently, tRFs have emerged as a distinct class of regulatory RNAs with potential immunoregulatory functions and altered tRF expression has been described during mycobacterial infection and active TB^18–20^. However, the longitudinal behavior of these molecules during treatment and their relationship to broader host transcriptional remodeling have not been well defined. In addition, interpretation of whole-blood transcriptomes is complicated by treatment-associated changes in circulating immune cell composition, making it difficult to distinguish changes in immune cell composition from cell-intrinsic transcriptional reprogramming^21^.

Here, we performed an integrative multi-omics analysis of paired whole-blood samples collected before and after four weeks of standard anti-tuberculosis therapy across three South African cohorts. By combining transcriptomic meta-analysis, immune deconvolution, and profiling of miRNAs and tRFs, we sought to define the coordinated host response during early treatment. We further developed a parsimonious baseline transcriptional signature, EARLY5, to investigate its association with early sputum culture conversion and assess its predictive contribution beyond baseline bacterial burden.

## Materials and Methods

### Study Design and Cohorts

This study used an integrative longitudinal multi-omics approach to characterize the host-response dynamics during early tuberculosis (TB) treatment. Three South African cohorts were included for discovery, integration, and validation (Figure 1): two publicly available transcriptomic datasets and a prospective longitudinal cohort recruited through the Regional Prospective Observational Research for Tuberculosis (RePORT-TB) consortium.

**Figure 1.**
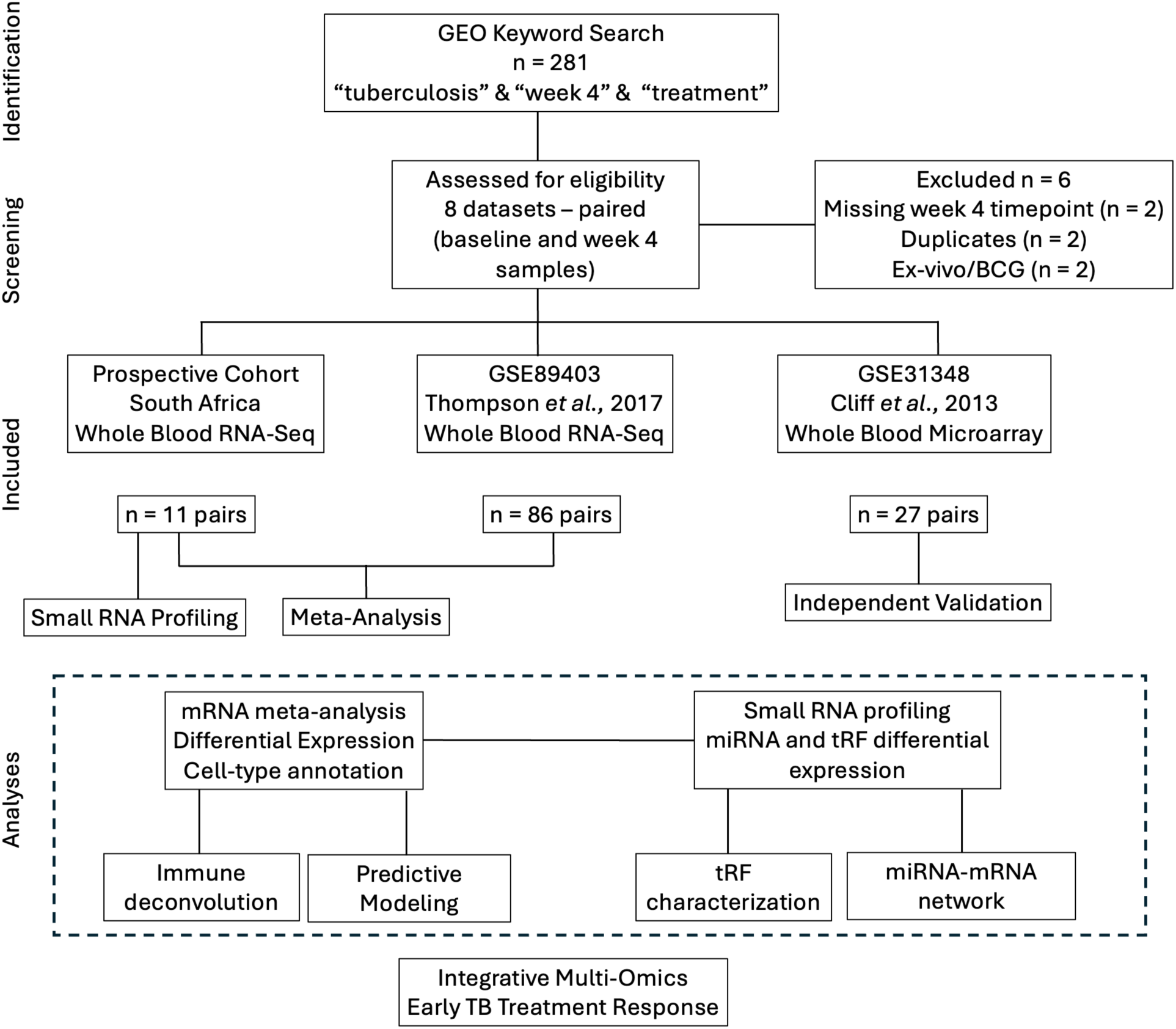
Study design and dataset selection flow. A systematic GEO search identified 281 tuberculosis treatment response datasets. Following initial screening, eight datasets underwent detailed eligibility assessment, of which six were excluded (missing 28-day timepoint, n = 2; duplicates, n = 2; ex vivo/BCG studies, n = 2). Two GEO datasets were retained: GSE89403 (n = 86 pairs; RNA-seq) and GSE31348 (n = 27 pairs; microarray). Together with the prospective South African cohort (n = 11 pairs; RNA-seq), three cohorts were included. The prospective cohort and GSE89403 were combined using Stouffer’s weighted Z-score meta-analysis for differential expression, while GSE31348 was used to assess directional concordance of treatment-associated transcriptional changes in an independent cohort. Downstream analyses included immune cell annotation, computational deconvolution, ncRNA profiling, miRNA–mRNA network analysis, tRF characterization, and predictive modeling.

The discovery and development cohort, GSE89403, is a whole-blood RNA sequencing dataset from South African adults with microbiologically confirmed pulmonary TB receiving standard first-line therapy^5^. 86 participants with paired baseline and week-4 samples were included in the meta-analysis. Of these, 76 had sputum culture conversion data and a baseline RNA-seq sample and were used for predictive modeling.

The independent validation cohort, GSE31348, comprised paired whole-blood microarrays from 27 South African adults with pulmonary TB, sampled at diagnosis and one month after treatment initiation^10^. It was used to assess reproducibility of treatment-associated transcriptional changes across platforms.

The prospective cohort comprised 11 participants with microbiologically confirmed drug-susceptible pulmonary TB recruited through the Perinatal HIV Research Unit, University of the Witwatersrand, Johannesburg, South Africa, under Institutional Review Board-approved RePORT-TB protocols. Paired whole-blood samples collected at treatment initiation and after four weeks of therapy were used for whole-transcriptome and small RNA sequencing. Clinical metadata, including sputum culture conversion status and baseline mycobacterial growth indicator tube (MGIT) time-to-positivity (TTP), were collected when available.

### Sample Collection and RNA Extraction

Peripheral whole blood (10 mL) was collected into PAXgene Blood RNA tubes (Qiagen) at baseline and after four weeks of treatment. Total RNA was isolated using the PAXgene Blood RNA Kit (Qiagen) according to the manufacturer’s instructions. RNA quantity and integrity were assessed before library preparation.

### Sequencing and Preprocessing

Library preparation and sequencing were performed by the Single Cell & Transcriptomics Core within the Johns Hopkins Integrated Genomics Center (Baltimore, MD, USA). For whole-transcriptome profiling, 500 ng of total RNA underwent poly(A) selection and was sequenced on an Illumina NovaSeq 6000. For small RNA profiling, 1 μg of RNA underwent size selection for 15-40 nucleotide fragments and was sequenced on an Illumina NextSeq 2000. Read quality was assessed using FastQC (v0.11.9) and MultiQC. Reads were aligned to the GRCh38.p14 human reference genome (GENCODE v47 annotation) using STAR (v2.7.10a) and gene-level counts were generated using featureCounts (Subread v2.0.3). Technical replicate counts in GSE89403 were summed before analysis. Raw CEL files from GSE31348 were normalized using robust multi-array average (RMA); probes were mapped using GPL570 annotations, and replicate probes were averaged before conversion to HGNC symbols using org.Hs.eg.db.

### Differential Expression and Meta-Analysis

Genes with low expression were filtered out (CPM > 1 in at least 10 samples in GSE89403, and at least 3 samples in the prospective cohort). RNA-seq data were normalized using the trimmed mean of M-values method (edgeR), voom-transformed, and analyzed using limma with paired designs and empirical Bayes moderation. Ensembl identifiers were mapped to HGNC symbols using Ensembl BioMart (release 116), genes with missing symbols were removed, and for duplicated symbols, the most highly expressed identifier was retained. GSE31348 was analyzed using limma with a paired design.

Results from GSE89403 and the prospective cohort were integrated using Stouffer’s weighted Z-score method. Gene-level p values were converted to signed Z scores according to fold-change direction and combined using weights proportional to the square root of cohort size, and combined p values were adjusted using the Benjamini–Hochberg procedure. Only genes detected in both cohorts with concordant direction were retained. Meta-differentially expressed genes (meta-DEGs) were defined by false discovery rate (FDR) < 0.05 and |log₂ fold change (FC)| ≥ 1, with meta-log₂FC calculated as the weighted average of cohort-specific effects. Directional concordance of meta-DEGs in GSE31348 was assessed using a binomial sign test. Each meta-DEG was assigned to the immune cell type with the highest normalized expression (nTPM) in the Human Protein Atlas. Genes with low expression across all HPA immune cells were assigned using published literature^4,22^. Reactome pathway enrichment was performed using Enrichr.

### Immune Cell Deconvolution

Immune cell composition in GSE89403 was estimated using CIBERSORTx (LM22, 1000 permutations) and MCP-counter^23–25^, using salmon-derived transcript-per-million (TPM) values^26^. Samples with CIBERSORTx-p < 0.05 were retained. Changes between baseline and week 4 were assessed using paired Wilcoxon signed-rank tests with FDR correction, with MCP-counter serving as orthogonal validation.

### Small RNA analysis

miRNAs and tRFs in the prospective cohort were quantified using miRge3.0 for read processing and annotation ^27,28^. tRNA-derived fragments (tRFs) were quantified using the cluster reference described by Lu et al.^27^, which groups reads by amino acid and anticodon to account for redundancy among tRNA isoacceptors; biogenesis class and mitochondrial origin were assigned from cluster annotations. Differential expression was assessed using DESeq2 with paired designs and apeglm shrinkage of log₂FC^29,30^. Features with ≥10 reads in at least 11 samples were tested, features with a base mean ≤ 50 were excluded, and differential expression was defined as FDR < 0.05 and |log₂FC| ≥ 0.58.

miRNA-mRNA interactions were identified using the multiMiR database, combining experimentally validated interactions (miRecords, miRTarBase, TarBase) together with predicted interactions (DIANA-microT, ElMMo, MicroCosm, miRanda, miRDB, PicTar, PITA, TargetScan) ^31^. miRNA-mRNA pairs in which the miRNA and its target changed in opposite directions were defined as inverse pairs, the pattern expected if the miRNA represses its target.

For read-level analyses of tRFs, reads from the parent tRNAs of each cluster were restricted to the cluster’s fragment type (for example, 3’-tRF reads for tRF-3 clusters). Seven-mer seed matching was performed, where the dominant sequence of each cluster was defined as its most abundant read, and reverse complements of nucleotides 2–8 were matched to the 3’ untranslated regions of meta-DEGs using TxDb and BSgenome annotations^32^. Anticodon-level changes were calculated as log₂ of the ratio of reads per 100,000 (RP100K) at week 4 to baseline, for anticodons contributing more than 5% of a cluster’s reads.

### RT-qPCR Validation

miR-144-3p, miR-144-5p and miR-32-5p were selected for validation based on effect size and biological relevance. They were measured in paired baseline and week-4 samples from eight participants in the prospective cohort. Total RNA (20 ng) was reverse transcribed using the miRCURY LNA RT Kit (Qiagen) with UniSp6 spike-in controls, and qPCR was performed using the miRCURY LNA SYBR Green PCR Kit (Qiagen) on a QuantStudio instrument. miR-103a-3p served as the endogenous reference for normalization, as its C_t_ values remained stable between baseline and week-4 samples. Samples with C_t_ > 36 or undetermined amplification were excluded. Relative expression was calculated using the 2^-ΔΔC^_t_ method, and baseline and week-4 ΔC_t_ values were compared using paired t-tests.

### Prediction of Early Sputum Culture Conversion

Participants in GSE89403 who were culture-negative by week 4 were classified as fast converters (n = 18) and the remainder as slow converters (n = 58). All predictive analyses used baseline samples. Gene pairs were selected using the k-top scoring pairs (k-TSP) algorithm (switchBox v1.44.0; up to five pairs), with the entire selection procedure nested within leave-one-out cross-validation (LOOCV). The out-of-fold EARLY5 score was the proportion of gene-pair votes for fast conversion (range 0–1). Because gene pairs were reselected within each fold, the cross-validated AUC estimates the performance of the full selection procedure rather than of a fixed set of pairs; the five pairs selected most frequently across folds are reported as the EARLY5 signature. Participants were classified as predicted fast converters when more than half of the gene-pair votes favored fast conversion.

Previously described Sweeney3 and RISK6 signatures were included as benchmark comparators^8,9^. Signature scores were calculated using TBSignatureProfiler (v1.20.0; GSVA v2.2.1)^33^. Discrimination was assessed using receiver operating characteristic (ROC) curves, with DeLong 95% CIs for the area under the curve (AUC), and paired DeLong tests were used to compare signatures (pROC v1.19.1). To assess the contribution of bacterial burden, EARLY5 was correlated with baseline MGIT TTP and Xpert Ct (Spearman), and logistic regression models of week-4 conversion including EARLY5, MGIT TTP or both were compared using a likelihood ratio test (p values for individual predictors were also obtained from likelihood ratio tests), with Firth regression (logistf v1.26.1) as a sensitivity analysis. Time to conversion was analyzed using Kaplan– Meier estimates and Cox proportional hazards models (survival v3.8-12), using Efron’s method for ties, with exact handling of ties as a sensitivity analysis.

### Statistical Analysis

All analyses were performed in R (version 4.5.0). Unless otherwise stated, paired and unpaired comparisons used Wilcoxon signed-rank and rank-sum tests, respectively, and correlations were assessed using Spearman’s rank correlation. For differential expression, deconvolution and multiple pairwise comparisons, p values were adjusted using the Benjamini-Hochberg procedure, with FDR < 0.05 considered significant. For all other analyses, two-sided p < 0.05 was considered statistically significant.

## Results

### Cohort characteristics

Three South African cohorts of adults with pulmonary tuberculosis were included in this study (Table 1). The prospective cohort comprised 11 participants with paired baseline and week 4 whole-blood RNA-seq samples. Median age was 35.5 years (IQR 29.3–51.5), and 10 participants (90.9%) were male. Three participants (27.3%) were HIV-positive, five (45.5%) were HIV-negative, and HIV status was unavailable for three (27.3%). Baseline AFB smear microscopy was positive in four participants (36.4%), and nine participants (81.8%) had positive baseline MGIT cultures with a median time-to-positivity of 9 days (IQR 3–11).

**Table 1.** Baseline characteristics of the study cohorts. Clinical characteristics of the prospective cohort, the GSE89403 cohort (used for meta-analysis and EARLY5 development) and the independent GSE31348 validation cohort. Paired baseline and week-4 samples were used for differential expression analyses in all cohorts. EARLY5 prediction analyses used baseline samples from GSE89403 only. Values are n (%) unless otherwise indicated. NR, not reported; TTP, time-to-positivity; Ct, cycle threshold. † Median (IQR); available for 10 of 11 participants (range 26–55 years). ᵃ 86 GSE89403 participants with paired baseline and week-4 samples were included in the meta-analysis. Of these, 76 had culture-conversion data and were used for EARLY5 development. The MGIT TTP, Xpert Ct, culture-conversion and treatment-outcome rows refer to these 76 participants.

| Characteristic | Prospective cohort<br>(n = 11) | GSE89403<sup>a</sup><br>(n = 86) | GSE31348<br>(n = 27) |
| --- | --- | --- | --- |
| Transcriptomic platform | RNA-seq | RNA-seq | Microarray |
| Samples analyzed | Baseline + week 4 (paired) | Baseline + week 4 (paired, n = 86) for meta-analysis; baseline only (n = 76) for prediction | Baseline + week 4 (paired) |
| Age, years | 35.5 (29.3–51.5)† | Adults 18–65 | Adults 18–65 |
| Male sex, n (%) | 10 (90.9) | NR | NR |
| Female sex, n (%) | 1 (9.1) | NR | NR |
| HIV-positive, n (%) | 3 (27.3) | 0 | 0 |
| HIV-negative, n (%) | 5 (45.5) | 86 (100) | 27 (100) |
| HIV status unknown, n (%) | 3 (27.3) | 0 | 0 |
| Pulmonary TB, n (%) | 11 (100) | 86 (100) | 27 (100) |
| Smear-positive TB, n (%) | 4 (36.4) | NR | 27 (100) |
| Culture-confirmed TB, n (%) | 9 (81.8) | 86 (100) | 27 (100) |
| Baseline MGIT TTP available, n (%) | 9 (81.8) | 58 (76.3) | NR |
| Baseline MGIT TTP, days, median (IQR) | 9 (3–11) | 5 (4–7) | NR |
| Baseline Xpert Ct available, n (%) | NR | 67 (88.2) | NR |
| Culture conversion by week 4, n (%) | — | 18 (23.7) | — |
| Treatment outcome | Cure (n = 2), relapse (n = 1), unknown (n = 8) | Definite cure (n = 67), probable cure (n = 6), possible cure (n = 3) | Cure |

GSE89403 comprised 86 HIV-negative adults with culture-confirmed pulmonary tuberculosis and paired baseline and week-4 whole-blood RNA-seq profiles, of whom 76 had culture-conversion data and a baseline sample for predictive modeling. The validation cohort, GSE31348, comprised 27 paired whole-blood microarrays from HIV-negative adults with smear-positive, drug-susceptible pulmonary tuberculosis who achieved durable cure.

### Meta-analysis identifies coordinated immune remodeling during early TB treatment

Stouffer’s weighted Z-score meta-analysis of GSE89403 and the prospective cohort identified 63 concordant meta-differentially expressed genes (meta-DEGs) between baseline and week 4 (FDR < 0.05, |log_2_FC| ≥ 1) (Supplementary Table S1; Figure 2a,b). Downregulated genes predominated (47/63, 75%) (Figure 2b), consistent with attenuation of inflammatory responses during early treatment.

**Figure 2.**
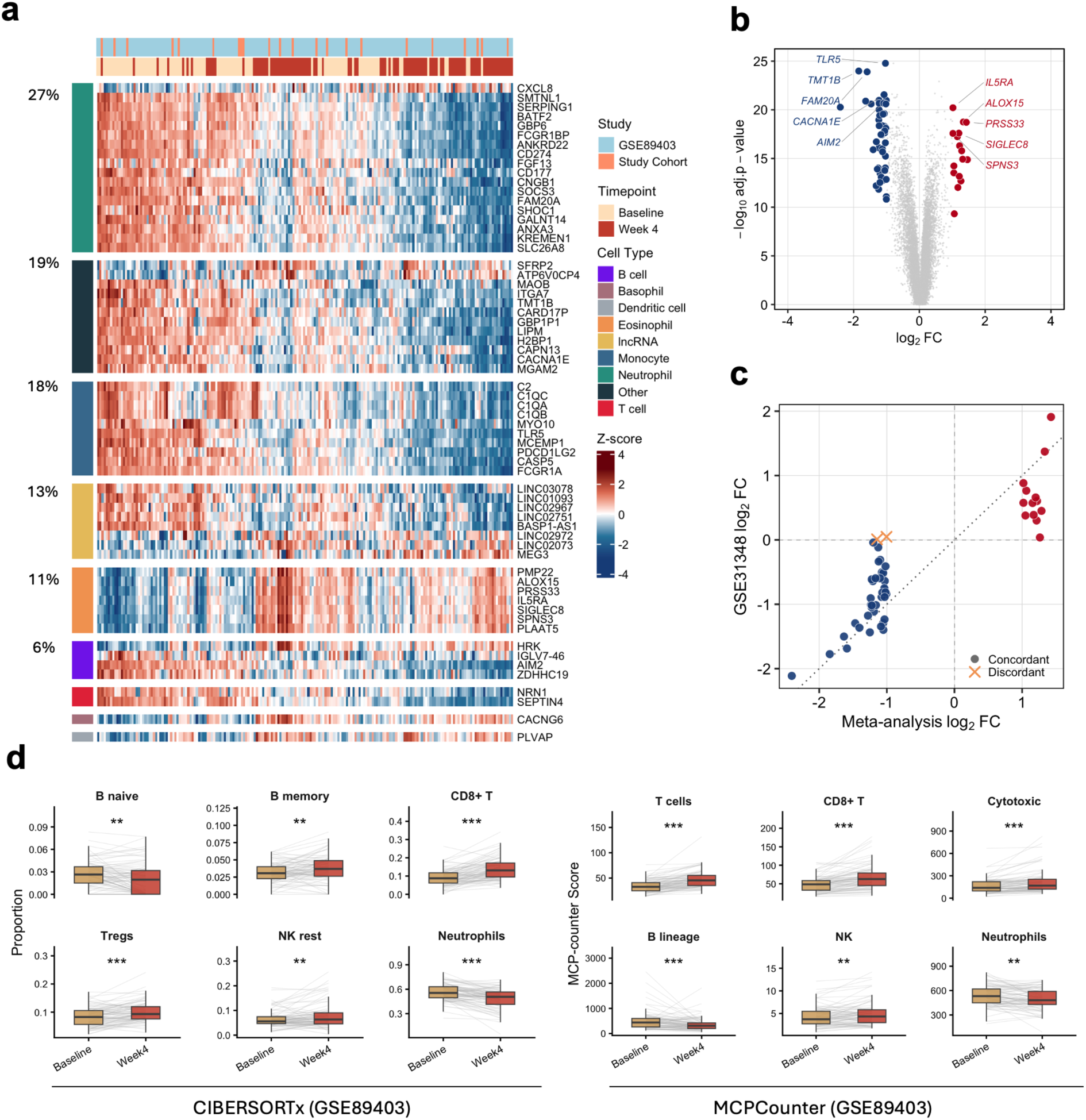
Meta-analysis identifies a coordinated innate-to-adaptive immune shift during early TB treatment. (a) Heatmap of 63 meta-DEGs across GSE89403 (n = 86 pairs) and the study cohort (n = 11 pairs), showing baseline-to-week 4 expression patterns with genes grouped by Human Protein Atlas cell-type annotation and hierarchically clustered within categories. (b) Volcano plot of Stouffer’s meta-analysis highlighting significantly up- and downregulated genes (FDR < 0.05, |log_2_FC| ≥ 1), with top hits labeled. (c) Validation in independent GSE31348 microarray data shows high directional consistency (51/53 genes, binomial test p = 3.2 × 10^−13^). (d) Paired immune deconvolution in GSE89403 demonstrates significant shifts in immune cell proportions and scores from baseline to week 4 (FDR < 0.05), consistent with treatment-associated changes in estimated innate and adaptive immune cell populations.

Cell-type annotation using Human Protein Atlas expression data showed that neutrophil-associated genes formed the largest category, including *CD177, BATF2, SERPING1, ANXA3, ANKRD22* and *CD274*, all of which were downregulated. Monocyte-associated genes were similarly suppressed, including complement components (*C1QA*, *C1QB*, *C1QC* and *C2*) and the immune checkpoint gene *PDCD1LG2*. In contrast, eosinophil-associated genes were upregulated, and the eight long non-coding RNAs among the meta-DEGs changed in both directions (Figure 2a). *TLR5* was the most significantly downregulated gene (log₂FC = −1.03, FDR = 1.68 × 10^−25^), *CD177* showed the largest reduction (log₂FC = −2.41), and *IL5RA* was the most significantly upregulated gene (log₂FC = 1.02, FDR = 6.12 × 10^−21^; Figure 2b).

In GSE31348, 51 of 53 meta-DEGs measured on the array changed in the same direction (binomial test p = 3.2 × 10^−13^), supporting reproducibility across independent cohorts and platforms (Figure 2c). Reactome pathway analysis showed enrichment of complement activation, innate immune signaling and interferon-γ signaling pathways (FDR < 0.05).

### Treatment-associated transcriptional changes are accompanied by shifts in estimated immune cell composition

Immune deconvolution of GSE89403 using CIBERSORTx and MCP-counter identified significant and concordant changes in circulating immune populations between baseline and week 4 (Figure 2d).

Neutrophil abundance decreased with both methods (FDR = 1.10 × 10^−7^ and 3.44 × 10^−3^, respectively), consistent with suppression of neutrophil-associated genes. In contrast, T-cell populations increased, including CD3⁺ T cells (FDR = 6.60 × 10^−14^), CD8⁺ T cells (FDR = 8.91 × 10^−10^ and 1.76 × 10^−11^), cytotoxic lymphocytes (FDR = 1.43 × 10^−7^) and regulatory T cells (FDR = 6.97 × 10^−6^). NK cells also increased with both methods (FDR = 8.63 × 10^−3^ and 3.44 × 10^−3^). B cells shifted from naive to memory phenotypes, with decreased naive B cells (FDR = 4.98 × 10^−3^) and increased memory B cells (FDR = 8.63 × 10^−3^), and total B-lineage abundance decreased (FDR = 4.71 × 10^−5^).

Monocyte and myeloid dendritic cell abundance did not change (FDR ≥ 0.18 with both methods; Supplementary Table S8), despite marked downregulation of monocyte-associated genes including *C1QA, C1QB, C1QC* and *PDCD1LG2*. These findings suggest that treatment-associated transcriptional changes may not be fully explained by estimated shifts in major immune cell populations, although cell-intrinsic effects cannot be established from bulk transcriptomic data.

### A baseline whole-blood signature is associated with early sputum culture conversion

Among 76 evaluable participants in GSE89403, sputum culture conversion occurred cumulatively in 18 (23.7%), 51 (67.1%), 68 (89.5%), and 76 (100%) participants by weeks 4, 8, 12, and 24, respectively (Figure 3a). Treatment outcome was classified as definite cure in 67 participants, probable cure in six and possible cure in three.

**Figure 3.**
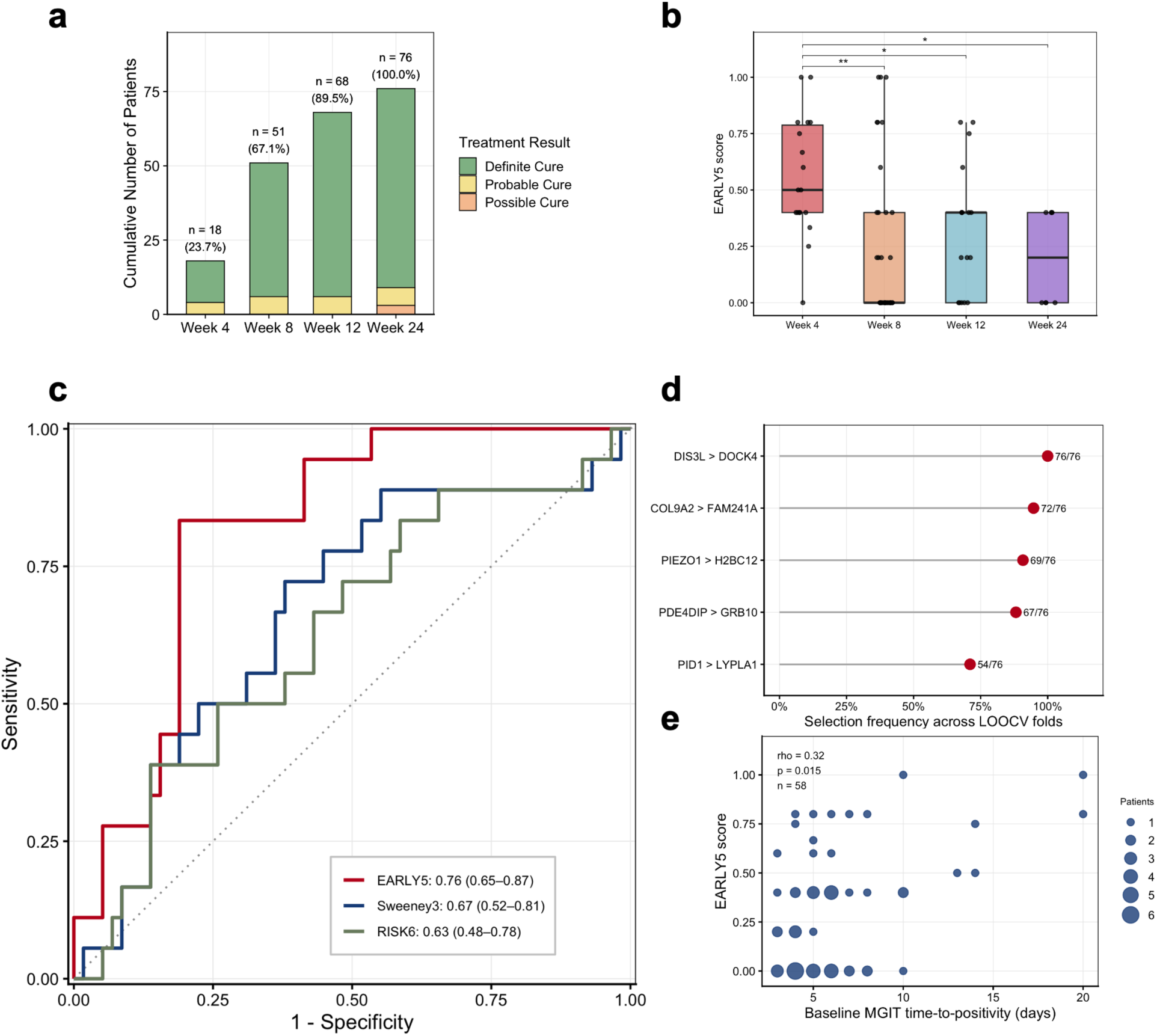
Baseline whole-blood EARLY5 score is associated with early sputum culture conversion in GSE89403. (a) Cumulative sputum culture conversion across weeks 4–24 (n = 76), colored by treatment outcome. (b) Baseline EARLY5 scores, calculated by leave-one-out cross-validation (LOOCV), were higher in participants who converted by week 4 than in those who converted later (Wilcoxon rank-sum test, BH-adjusted; *p < 0.05, **p < 0.01). (c) ROC curves for predicting week-4 conversion. EARLY5 (AUC 0.76, 95% CI 0.65–0.87) showed numerically higher discrimination than Sweeney3 (0.67, 0.52–0.81) and RISK6 (0.63, 0.48–0.78), although the differences were not statistically significant (paired DeLong test, p = 0.171 and 0.113). (d) Four of the five EARLY5 gene pairs were selected in at least 88% of the 76 LOOCV folds. (e) EARLY5 scores correlated modestly with baseline MGIT time-to-positivity (n = 58; Spearman rho = 0.32, p = 0.015), point size reflects the number of participants.

Across LOOCV folds, the five most frequently selected gene pairs were *DIS3L–DOCK4, COL9A2–FAM241A, PIEZO1–H2BC12, PDE4DIP–GRB10* and *PID1–LYPLA1*, which defined the final EARLY5 signature. Four of the five were selected in at least 88% of the 76 folds, and no other pair was selected in more than 7% (Figure 3d; Supplementary Table S7). Out-of-fold EARLY5 scores were higher in week-4 converters than in participants converting at week 8 (BH-adjusted p = 0.004), week 12 (p = 0.024) or week 24 (p = 0.010; Figure 3b).

EARLY5 discriminated week-4 conversion with a cross-validated AUC of 0.76 (95% CI 0.65–0.87). The published Sweeney3 and RISK6 signatures had AUCs of 0.67 (0.52– 0.81) and 0.63 (0.48–0.78), respectively. These did not differ significantly from EARLY5 (paired DeLong p = 0.171 and 0.113; Figure 3c; Supplementary Table S5).

EARLY5 scores correlated modestly with baseline TTP (Spearman ρ = 0.32, p = 0.015; Figure 3e), with higher scores associated with lower bacterial burden, but not with Xpert C_t_ (ρ = 0.20, p = 0.114; Supplementary Figure S2). In the 58 participants with TTP data, EARLY5 and TTP were each associated with week-4 conversion (Table 2). In a combined model, EARLY5 remained associated with conversion (adjusted OR 1.86 per gene-pair vote, 95% CI 1.10–3.46, p = 0.021), and adding EARLY5 improved model fit over TTP alone (likelihood ratio test p = 0.021). Results were similar with Firth penalized regression (p = 0.026).

**Table 2.** Association of baseline EARLY5 score and bacterial burden (MGIT TTP) with week-4 sputum culture conversion. Logistic regression of culture conversion by week 4 (fast) vs later in GSE89403 participants with baseline MGIT time-to-positivity (TTP) available (n = 58; 12 fast converters). All models were fitted to the same 58 participants. The EARLY5 score is the out-of-fold LOOCV proportion of gene-pair votes for fast conversion (range 0-1); ORs are per 0.2 increase, equivalent to one gene-pair vote. MGIT TTP is expressed per interquartile range (IQR); longer TTP indicates lower baseline bacterial burden. 95% CIs are profile likelihood intervals. P values are from likelihood ratio tests.

| Model | Predictor | OR (95% CI) | p |
| --- | --- | --- | --- |
| EARLY5 only | EARLY5 score (per 0.2) | 2.26 (1.41–4.06) | <0.001 |
| MGIT TTP only | MGIT TTP (per IQR) | 2.56 (1.45–5.48) | <0.001 |
| EARLY5 + MGIT TTP | EARLY5 score (per 0.2) | 1.86 (1.10–3.46) | 0.021 |
|  | MGIT TTP (per IQR) | 2.00 (1.05–4.51) | 0.033 |
| Likelihood ratio test: EARLY5 + MGIT TTP vs MGIT TTP only | — | — | 0.021 |

In a time-to-event analysis, participants predicted to be fast converters converted earlier (log-rank p = 0.020; HR 1.85, 95% CI 1.08–3.16). In the 58 participants with MGIT TTP data (unadjusted HR 1.96, 95% CI 1.06–3.65), the association was attenuated after adjustment for MGIT TTP (HR 1.72, 0.90–3.31, p = 0.102; Supplementary Figure S1). Conclusions were unchanged with exact handling of ties (Supplementary Table S6).

### Whole-blood miRNA profiling identifies treatment-responsive species linked to the mRNA landscape

Differential expression analysis of small RNA-seq data from the prospective cohort identified seven miRNAs significantly altered between baseline and week 4 (FDR < 0.05, |log_2_FC| ≥ 0.58, base mean > 50) (Supplementary Table S2, Figure 4a,b). Three were upregulated and four downregulated (Figure 4a). miR-151a-5p/151b was the most significantly downregulated miRNA (log_2_FC = −0.64, FDR = 8.3 × 10^−5^), whereas miR-144-3p and miR-144-5p, two arms of the same precursor, were among the most strongly upregulated (log_2_FC = 1.09 and 0.94; FDR = 0.037 and 0.031, respectively) (Figure 4b).

**Figure 4.**
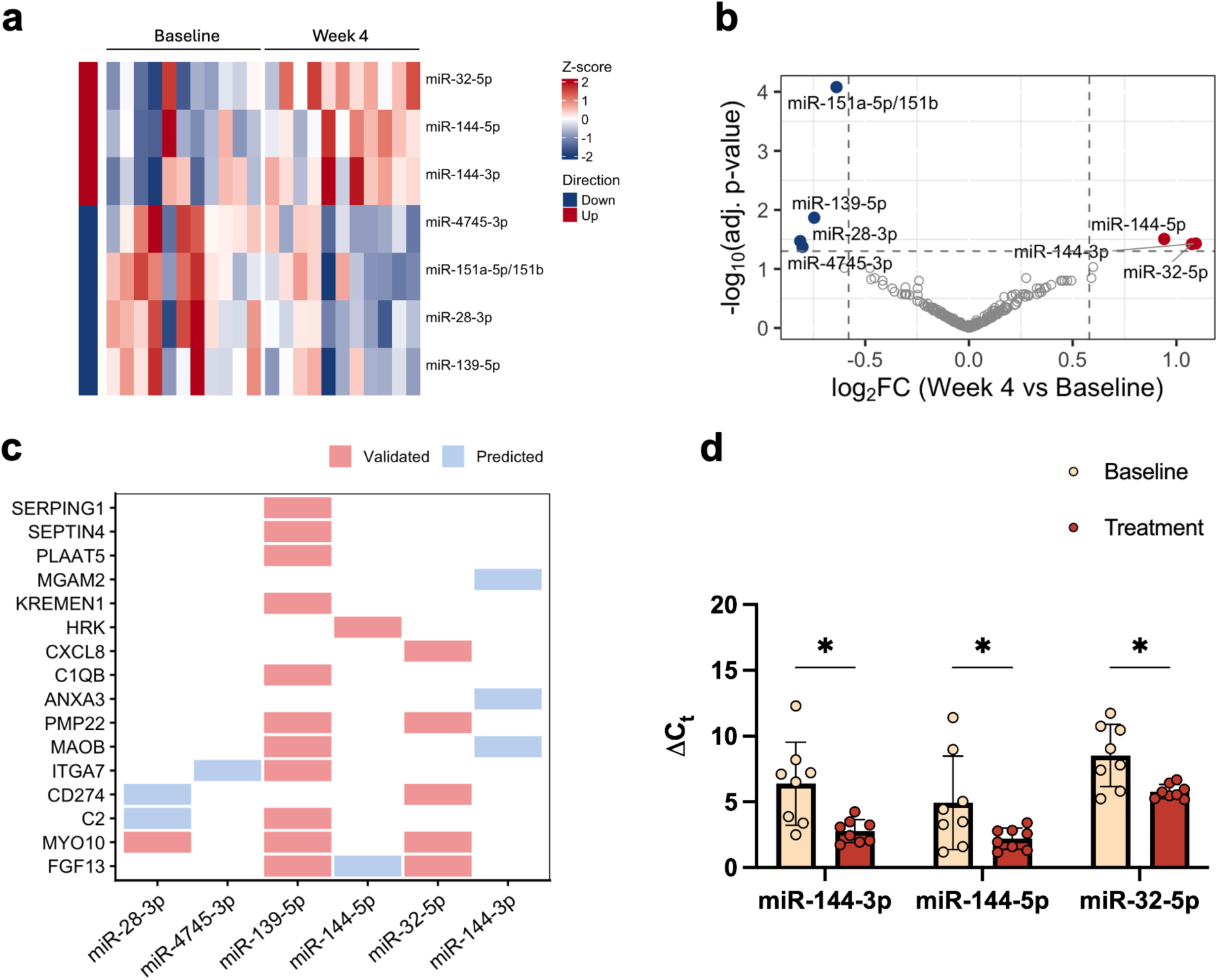
Differentially expressed miRNAs and their interactions with mRNA meta-DEGs during TB treatment. (a) Heatmap of 7 DE-miRNAs across paired baseline and week 4 samples (n = 11 pairs), showing consistent within-subject expression changes; rows annotated by directionality (up- vs downregulated). (b) Volcano plot of miRNA differential expression (n = 227; base mean > 50), highlighting significant changes (|log_2_FC| ≥ 0.58, FDR < 0.05; DESeq2 with apeglm shrinkage). (c) miRNA–mRNA interaction network linking DE-miRNAs to meta-DEGs, integrating validated and predicted targets (multiMiR). (d) RT-qPCR confirmation of three upregulated miRNAs in a subset of the same prospective cohort (n = 8 pairs), using miR-103a-3p as a stable endogenous reference (paired t-test, *p < 0.05).

Integration with the 63 meta-DEGs using multiMiR identified 25 miRNA-mRNA interactions involving six of the seven differentially expressed miRNAs, including 18 experimentally validated interactions (Supplementary Table S3, Figure 4c). miR-139-5p had the largest number of validated target interactions, including genes involved in complement activation (*C1QB, C2, SERPING1*). Upregulated miRNAs, including miR-32-5p and miR-144-3p, had validated or predicted interactions with several downregulated meta-DEGs, including *FGF13*, *ANXA3*, *CD274*, and *MYO10*.

Nine miRNA-mRNA pairs showed inverse regulation (Supplementary Table S3). RT-qPCR analysis of eight paired samples from the prospective cohort confirmed significant upregulation of miR-144-3p, miR-144-5p, and miR-32-5p at week 4 compared with baseline (miR-144-3p, p = 0.0099; miR-144-5p, p = 0.045; miR-32-5p, p = 0.015, paired t-test) (Figure 4d).

### tRNA-derived fragments showed treatment-responsive, isoacceptor-specific cleavage patterns

Eight tRFs changed between baseline and week 4 (FDR < 0.05, |log_2_FC| ≥ 0.58) (Supplementary Table S4, Figure 5a, b), seven upregulated and one downregulated fragment, Asp Comb 3 (log2FC = −0.65, FDR = 0.034). Gly Comb 2 (log_2_FC = 1.02, FDR = 0.0024) and Ala Comb 2 (log_2_FC = 0.97, FDR = 0.0078) were the most significantly upregulated (Figure 5b).

**Figure 5.**
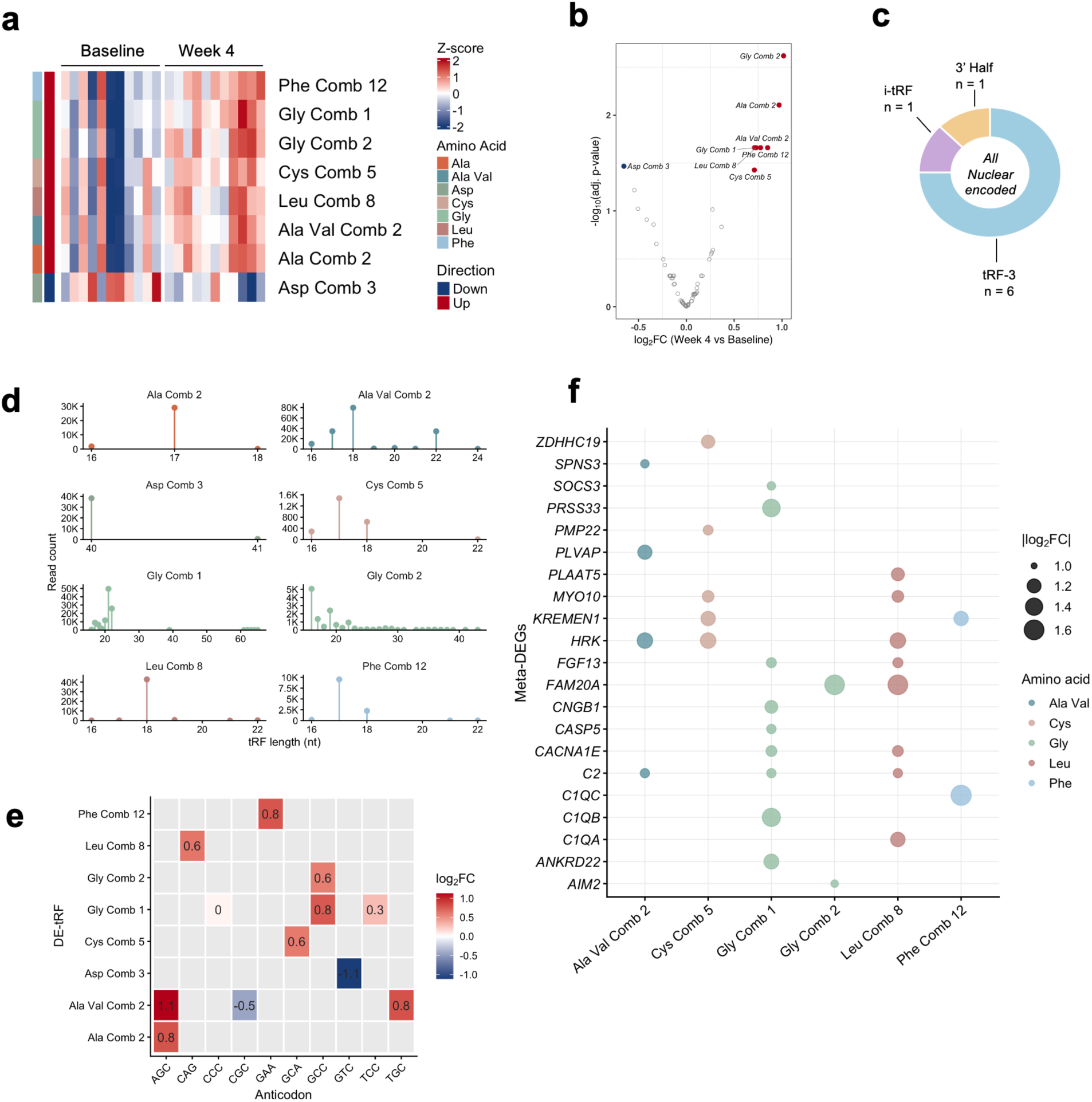
Eight nuclear-encoded tRNA-derived fragments are differentially expressed during early tuberculosis treatment. (a) Heatmap of Z-scored expression for the 8 DE-tRFs across paired baseline and week-4 samples (n = 11 pairs) with columns grouped by timepoint and rows annotated by amino acid family and directionality. (b) Volcano plot of tRF differential expression (DESeq2 with apeglm shrinkage; significance thresholds: FDR < 0.05, |log_2_FC| ≥ 0.58); significant tRFs labeled and colored by direction. (c) Biogenesis classification of the 8 DE-tRFs by fragment type (tRF-3, n = 6; i-tRF, n = 1; 3’ half, n = 1); all are nuclear-encoded. (d) Read-length distributions for each DE-tRF, illustrating dominant fragment lengths and variation across tRF clusters. (e) Anticodon-level log_2_FC heatmap showing abundance changes within each DE-tRF cluster; values represent fold change per anticodon species, demonstrating within-cluster isoacceptor specificity. (f) Predicted tRF-mRNA interactions based on 7-mer seed matching to meta-DEG 3’ UTRs (TxDb/BSgenome hg38); dot size reflects absolute meta-analysis log_2_FC and color indicates tRF amino acid family.

All eight tRFs were nuclear-encoded, with no mitochondrial-derived fragments identified. Biogenesis analysis demonstrated a bias toward tRF-3 fragments (6/8). Remaining species included one internal tRF (Gly Comb 2) and one 3’ half fragment (Asp Comb 3) (Figure 5c). Length distribution analysis revealed dominant fragment lengths for most DE-tRFs (17–21 nt for tRF-3 species and 40 nt for Asp Comb 3), with broader distributions for Gly Comb 1 and Gly Comb 2, indicating distinct fragment-length profiles across tRF clusters (Figure 5d).

Anticodon-level analysis revealed isoacceptor bias within DE-tRF clusters (Figure 5e). Of the four clusters containing more than one isoacceptor, three were dominated by a single anticodon: Gly Comb 1 and Gly Comb 2 were driven predominantly by GCC (log₂FC = 0.8 and 0.6), with a minor contribution from TCC in Gly Comb 1 (log₂FC = 0.3), and Leu Comb 8 by CAG (log₂FC = 0.6). Ala Val Comb 2 showed the broadest anticodon distribution, with increases in AGC (log₂FC = 1.1) and TGC (log₂FC = 0.8) but a decrease in CGC (log₂FC = −0.5). The four single-isoacceptor clusters changed in the same direction as their overall abundance: Phe Comb 12 (GAA, log₂FC = 0.8), Cys Comb 5 (GCA, log₂FC = 0.6) and Ala Comb 2 (AGC, log₂FC = 0.8) increased, and Asp Comb 3 (GTC, log₂FC = −1.1) decreased. These findings suggest isoacceptor-associated differences in tRF abundance during treatment, although the underlying mechanisms remain unclear.

Seven-mer seed matching against the 3’UTRs of the 63 meta-DEGs identified 30 putative tRF-mRNA interactions across six of the eight DE-tRFs (Figure 5f). Gly Comb 1 and Leu Comb 8 had the broadest predicted target coverage (nine and eight genes, respectively). Predicted targets included the complement genes *C1QB* (Gly Comb 1), C1QA (Leu Comb 8), *C1QC* (Phe Comb 12) and *C2* (Gly Comb 1, Leu Comb 8 and Ala Val Comb 2). All six tRFs with predicted targets were upregulated, and 22 of the 30 predicted targets were downregulated, similar to the proportion of downregulated genes among all meta-DEGs (47/63).

## Discussion

This study presents an integrative multi-omics characterization of whole-blood transcriptional dynamics during early tuberculosis (TB) treatment, combining mRNA meta-analysis from two South African cohorts, directional validation in a third cohort, and paired miRNA and tRF profiling from a prospective longitudinal cohort. The findings indicate attenuation of innate inflammatory programs and complement signaling, shifts in estimated immune cell populations, and increased expression of erythroid-associated miRNAs during early treatment, alongside identification of a baseline whole-blood signature, EARLY5, associated with early sputum culture conversion.

The mRNA meta-analysis identified 63 genes consistently dysregulated between baseline and week 4, with downregulation predominating and concentrated among neutrophil- and monocyte-associated genes. These findings are consistent with prior descriptions of interferon-inducible, neutrophil-driven blood transcriptional signatures during active TB and their contraction during treatment^4,10^. Immune deconvolution findings were consistent with Human Protein Atlas-based cell type annotations, indicating reduced estimated neutrophil abundance and increased estimated representation of adaptive immune populations, including CD8+ T cells. Notably, estimated monocyte abundance remained relatively stable despite marked suppression of monocyte-associated transcripts, suggesting that changes in transcriptional activity may occur without corresponding changes in estimated monocyte abundance. These findings are consistent with coordinated immune remodeling during early bacterial clearance.

Three major transcriptional programs emerged from the meta-DEG landscape. The first was broad contraction of innate inflammatory signaling, including suppression of interferon-stimulated genes, inflammasome-associated genes, and complement pathway components including *C1QA, C1QB, C1QC, C2,* and *SERPING1*. *SERPING1* and *SEPTIN4* have been identified previously as markers of active TB, reinforcing that their downregulation reflects a transition away from active disease^34^. Complement activation is a well-recognized feature of active TB immunopathology, contributing both to opsonization and local immune dysregulation within granulomatous lesions^35^. The reduction in complement-associated transcripts by week 4 suggests that changes in complement-related transcription may occur during the early phase of tuberculosis treatment.

The second program reflected active immune resolution. Eosinophil-associated genes, including *IL5RA, SIGLEC8, PRSS33*, and the pro-resolving lipid mediator enzyme *ALOX15*, were uniformly upregulated, consistent with the increase in eosinophils estimated by CIBERSORTx (Supplementary Table S8). Three of eight long non-coding RNAs were also upregulated. The third program involved changes in adaptive immunity-associated transcriptional patterns. Downregulation of checkpoint genes *CD274* and *PDCD1LG2* occurred alongside increased estimated CD8+ T-cell abundance, suggesting treatment-associated changes in adaptive immune responses.

The EARLY5 classifier suggests that baseline whole-blood transcriptomic profiles contain information associated with early sputum culture conversion within the discovery cohort. Sweeney3 and RISK6 showed numerically lower performance that did not differ significantly from EARLY5, possibly reflecting their development for diagnosis and treatment monitoring rather than prediction of sputum conversion kinetics^8,9^. This comparison also favors EARLY5, which was developed for this outcome in this cohort, whereas Sweeney3 and RISK6 were applied without retraining. EARLY5 scores correlated modestly with baseline MGIT TTP, indicating partial overlap with bacterial burden. Although EARLY5 remained associated with week-4 conversion after adjustment for MGIT TTP in logistic regression, the small number of early converters and attenuation of the adjusted time-to-conversion association limit conclusions regarding its independent prognostic value. The wide confidence interval around the AUC also means that the comparison with published signatures was not powered to detect modest differences. These findings provide proof-of-concept that pretreatment blood transcriptional profiles may inform treatment response stratification in future studies.

The miRNA findings add a post-transcriptional layer to the mRNA landscape. The most significant DE-miRNA, miR-151a-5p/151b, had no identifiable targets among the 63 meta-DEGs, suggesting involvement in pathways not captured by the present mRNA signature. The miR-144/451 locus, regulated by GATA1 and essential for erythroid homeostasis, emerged as a prominent feature of treatment response. miR-144-3p and miR-144-5p are among the most abundant miRNAs in mature erythrocytes^36^. Their increase at week 4 may reflect changes in erythroid cell composition or erythropoietic activity during treatment, although these mechanisms were not directly assessed^37^. This interpretation is consistent with the well-recognized anemia of chronic disease in active TB and with previous reports linking miR-144-3p to treatment response^14^. Although miR-144 can regulate autophagy in macrophage models via ATG4A and DRAM2^12^, the relevance of these mechanisms in whole blood, where the erythroid signals are likely dominant, remains unclear.

Within our dataset, among the inverse pairs, miR-32-5p, whose increase was confirmed by RT-qPCR, has experimentally supported interactions with CD274, FGF13 and MYO10, all of which were downregulated, and miR-144-3p is predicted to target the downregulated ANXA3, MAOB and MGAM2. These inverse expression patterns suggest potential miRNA-mRNA regulatory relationships, although direct repression cannot be established from the present data. In addition, miR-144 is largely erythroid, whereas targets such as ANXA3 are expressed mainly in neutrophils, and bulk blood measurements also reflect shifts in cell composition, so the inverse pattern may also arise without direct regulation. The increase in miR-32-5p is notable given its reported role in *Mtb*-infected macrophages, where TLR4-driven induction suppresses FSTL1 and dampens inflammatory cytokine responses^38^. Its increase during treatment may reflect context-dependent immune dampening during resolution, though prior findings are based on isolated macrophage models and may not directly translate to whole blood.

The tRF findings characterize the cellular tRF landscape during TB treatment in patient-derived whole blood, a context distinct from prior work examining extracellular circulating tRFs as diagnostic biomarkers in active infection and intracellular tRF dysregulation in isolated macrophages infected in vitro^18,19^. Notably, the in vitro macrophage models involved direct *Mtb* infection at relatively high bacillary burdens, whereas circulating cells in PAXgene whole blood samples are largely uninfected and respond primarily to systemic inflammatory signals. Mitochondrial tRFs may therefore reflect infection-associated cellular stress in directly infected cells, whereas nuclear-encoded tRFs identified during treatment may instead reflect restoration of cellular homeostasis^18^. The predominance of 17–21 nt tRF-3 fragments is notable, as these species are generated via Dicer- and angiogenin-independent pathways and can associate with Argonaute complexes to mediate post-transcriptional gene silencing^39^.

Anticodon-level analysis revealed that tRF abundance changes were concentrated in one or two isoacceptors within each amino acid family rather than distributed uniformly, suggesting non-random, isoacceptor-specific regulation of tRF abundance. Although angiogenin preferentially targets specific tRNA isoacceptors in other systems, the predominance of tRF-3 species here suggests that the responsible nuclease(s) remain undefined^40^. Whether this anticodon specificity reflects selective biogenesis or differences in tRNA isoacceptor expression among the changing blood cell populations cannot be resolved from the current data. More broadly, because small RNAs were measured in whole blood, changes in miRNA and tRF abundance may partly reflect the shifts in neutrophil, lymphocyte and erythroid composition described above.

A notable feature of the integrated analysis is that complement genes, which were among the most prominently downregulated mRNAs during early treatment, were predicted targets of several upregulated tRFs: C1QB (Gly Comb 1), C1QA (Leu Comb 8), C1QC (Phe Comb 12) and C2 (three tRFs). Although these predicted interactions suggest a potential role for tRFs in complement transcript regulation, direct regulatory activity remains to be experimentally established. In contrast, miR-139-5p, which has validated interactions with C1QB, C2 and SERPING1, decreased together with these genes and is therefore unlikely to drive their suppression. The tRF-complement interactions are based on seed complementarity alone, and the downregulation of these genes may also reflect changes in monocyte transcription.

## Limitations

The prospective cohort was small, predominantly male (90.9%) and included participants living with HIV, unlike the HIV-negative public cohorts, for which sex was not reported. Small RNA findings come from 11 participants without independent sequencing replication, and RT-qPCR was performed in a subset of the same participants. Bulk profiling cannot fully resolve cell-type-specific effects, and the miRNA- and tRF-mRNA interactions have not been tested experimentally. EARLY5 was developed in a single cohort and requires external validation, and burden-adjusted analyses included only 58 participants (12 fast converters). Finally, peripheral blood may not fully reflect immune dynamics at the site of infection.

## Conclusions

This study provides an integrative multi-omics characterization of the early host response to tuberculosis treatment, revealing coordinated changes in innate and adaptive immune transcriptional programs by week 4 that may reflect both shifts in immune cell composition and changes in transcriptional activity. Small RNA profiling identified treatment-responsive miRNAs, including erythroid-associated miR-144, and a nuclear-encoded, tRF-3-dominated tRF response that differs from the mitochondria-biased pattern reported in *Mtb*-infected macrophages. Inverse miRNA-mRNA pairs and predicted tRF targets among downregulated complement genes suggest that post-transcriptional regulation may contribute to these changes. In addition, the baseline EARLY5 signature was associated with week-4 sputum culture conversion and remained associated after adjustment for MGIT TTP in a subset of participants, supporting its potential as an exploratory prognostic biomarker requiring independent validation. These findings provide a multi-omics framework for understanding early treatment response and for developing clinically translatable biomarkers.

## Ethics and IRB Statement

The prospective study was approved by the Johns Hopkins University Institutional Review Board and the University of the Witwatersrand Human Research Ethics Committee. Written informed consent was obtained from all participants. Publicly available datasets GSE89403 and GSE31348 were analyzed in accordance with their original ethical approvals, as described in the respective primary publications.

## Data and Code Availability

Sequencing data from the prospective study cohort have been deposited in the NCBI Gene Expression Omnibus (GEO) under accession number GSE341867. Publicly available datasets GSE89403 and GSE31348 are accessible via the NCBI Gene Expression Omnibus. Analysis code is available at https://github.com/AnoushkaGV/integrative-multi-omics-mtb.git

## Contributors

GVAC, DQ, and PCK conceived and designed the study. GVAC, DQ, JSB, MKH, ATLQ, and PCK developed the methodology. NM contributed to investigation and provided resources. GVAC and DQ curated the data and GVAC, DQ and ATLQ performed the analysis. GVAC and PCK wrote the original draft of the manuscript, and all authors reviewed and edited the manuscript. BBA, MKH, JSB, NM, and PCK administered the project. PCK acquired funding.

## Acknowledgements

We thank the study participants, the staff of the Perinatal HIV Research Unit (University of the Witwatersrand), and the RePORT South Africa consortium. We thank the Single Cell & Transcriptomics Core at the Johns Hopkins Integrated Genomics Center for sample processing and library preparation, and the investigators who generated the GSE89403 and GSE31348 datasets.

## Declaration of interests

We declare no competing interests.

## Supplementary Material

**Supplementary Figure S1.**
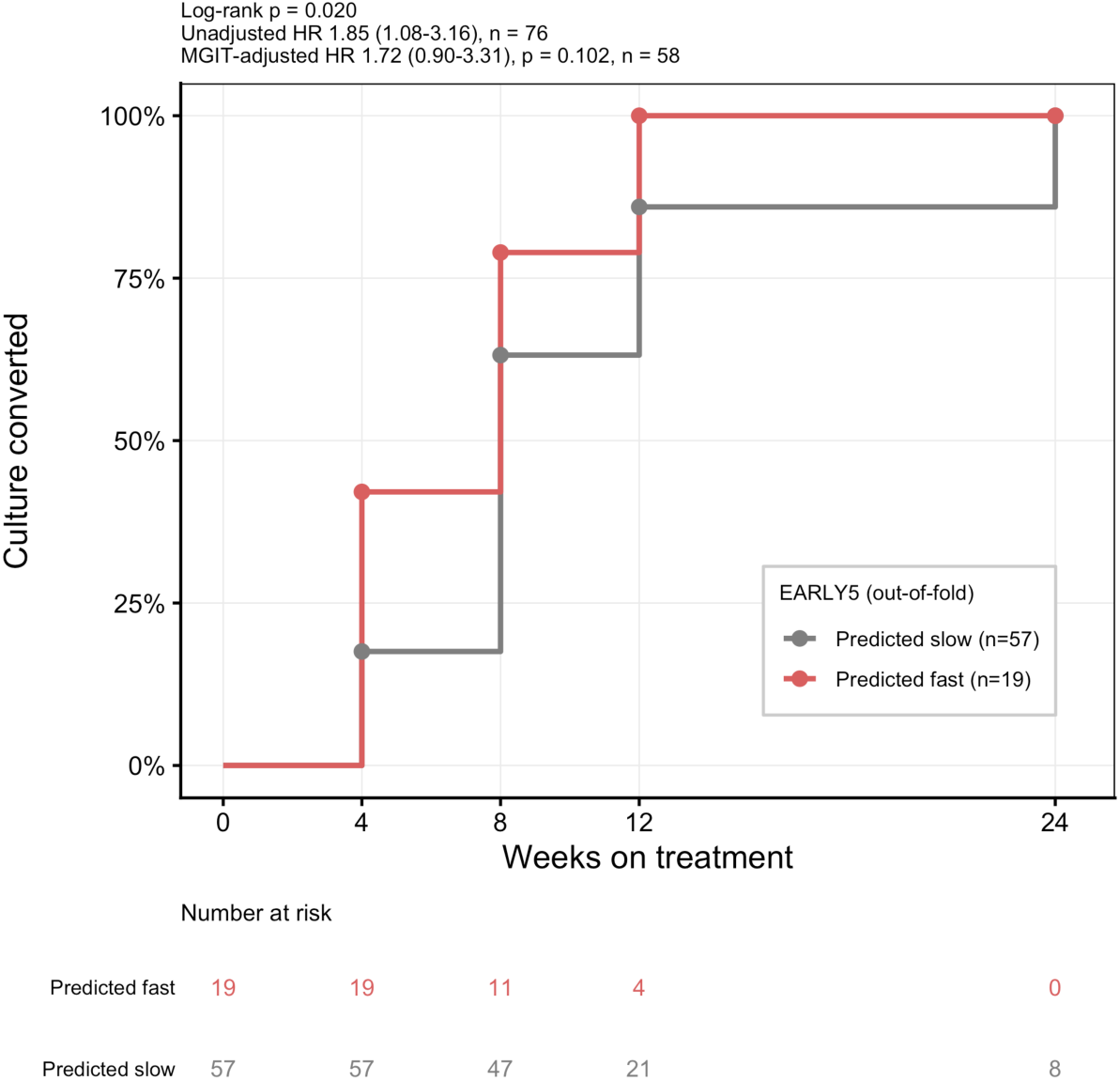
Time to sputum culture conversion by EARLY5 prediction. Cumulative proportion of participants with sputum culture conversion in GSE89403, grouped by out-of-fold EARLY5 classification (predicted fast, n = 19; predicted slow, n = 57). Participants predicted to convert fast did so earlier (log-rank p = 0.020; HR 1.85, 95% CI 1.08–3.16). After adjustment for baseline MGIT time-to-positivity, the association was attenuated (HR 1.72, 95% CI 0.90–3.31, p = 0.102; n = 58). Culture conversion was assessed at weeks 4, 8, 12 and 24. Numbers at risk are shown below the plot.

**Supplementary Figure S2.**
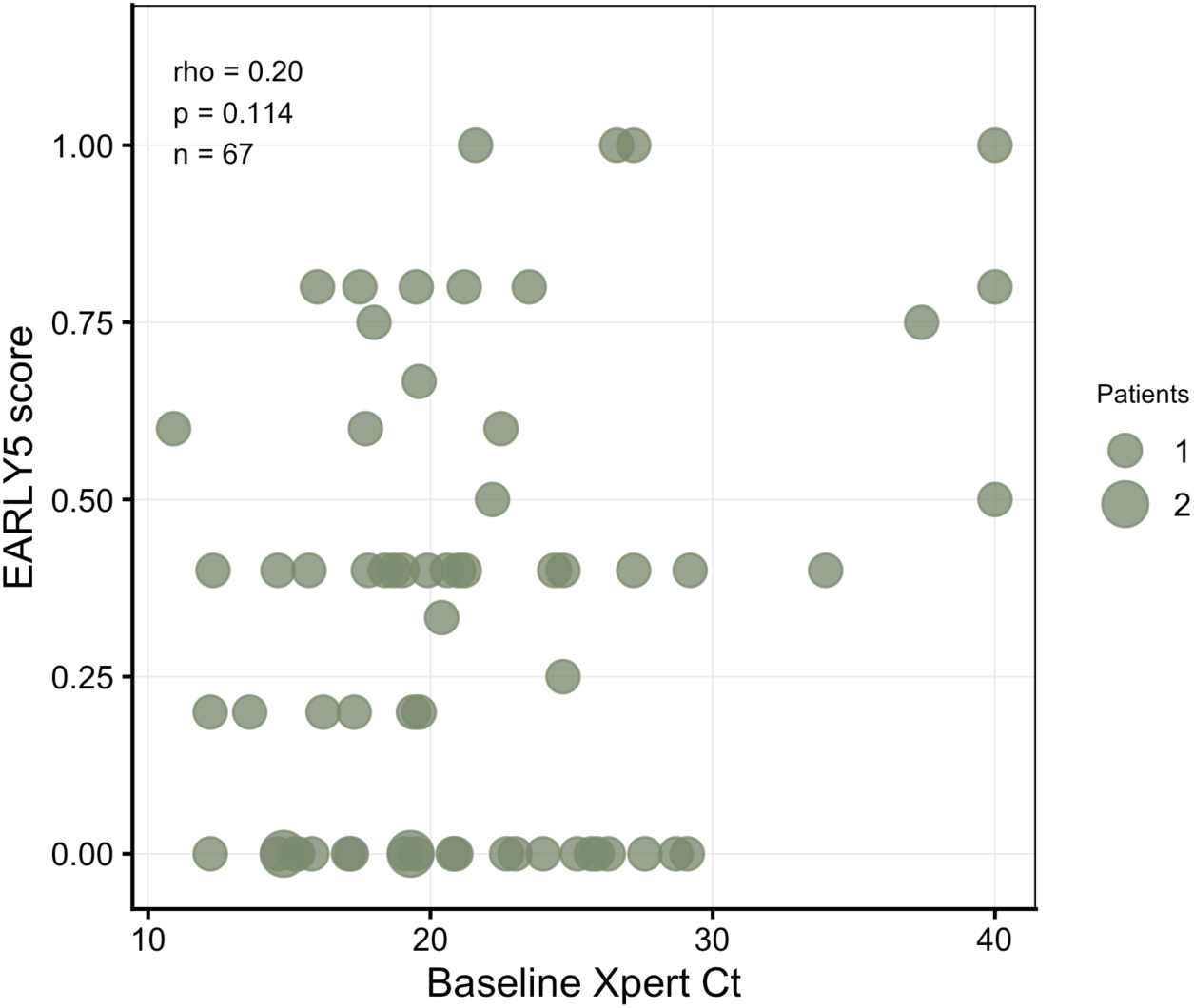
EARLY5 score and baseline Xpert Ct. EARLY5 LOOCV scores were not significantly correlated with baseline Xpert cycle threshold (n = 67; Spearman rho = 0.20, p = 0.114). Ct values of 40 indicate no detection (n = 3). Excluding these participants, the correlation was weaker (n = 64; Spearman rho = 0.10, p = 0.43). Point size reflects the number of participants at each value.

**Supplementary Table S1.**
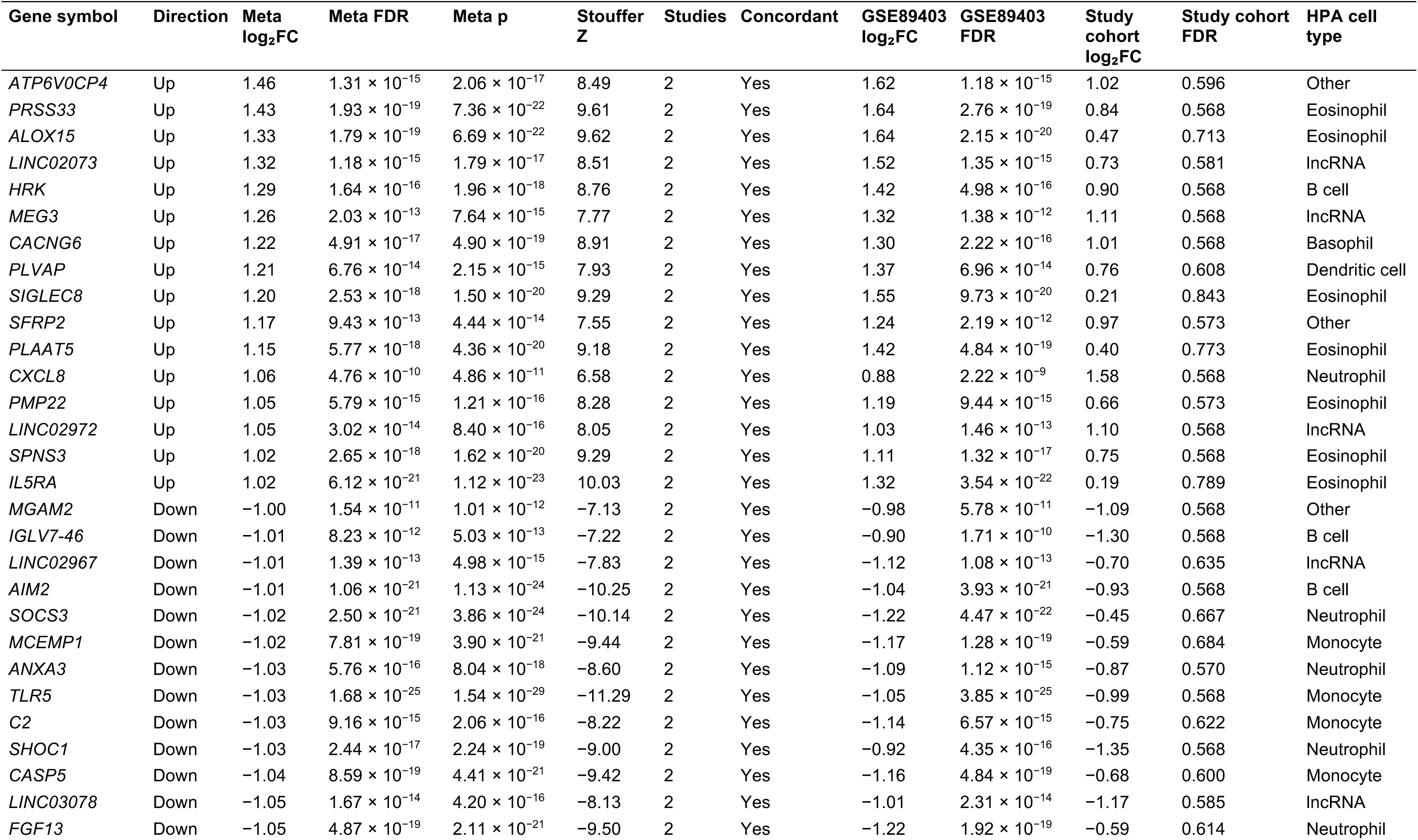
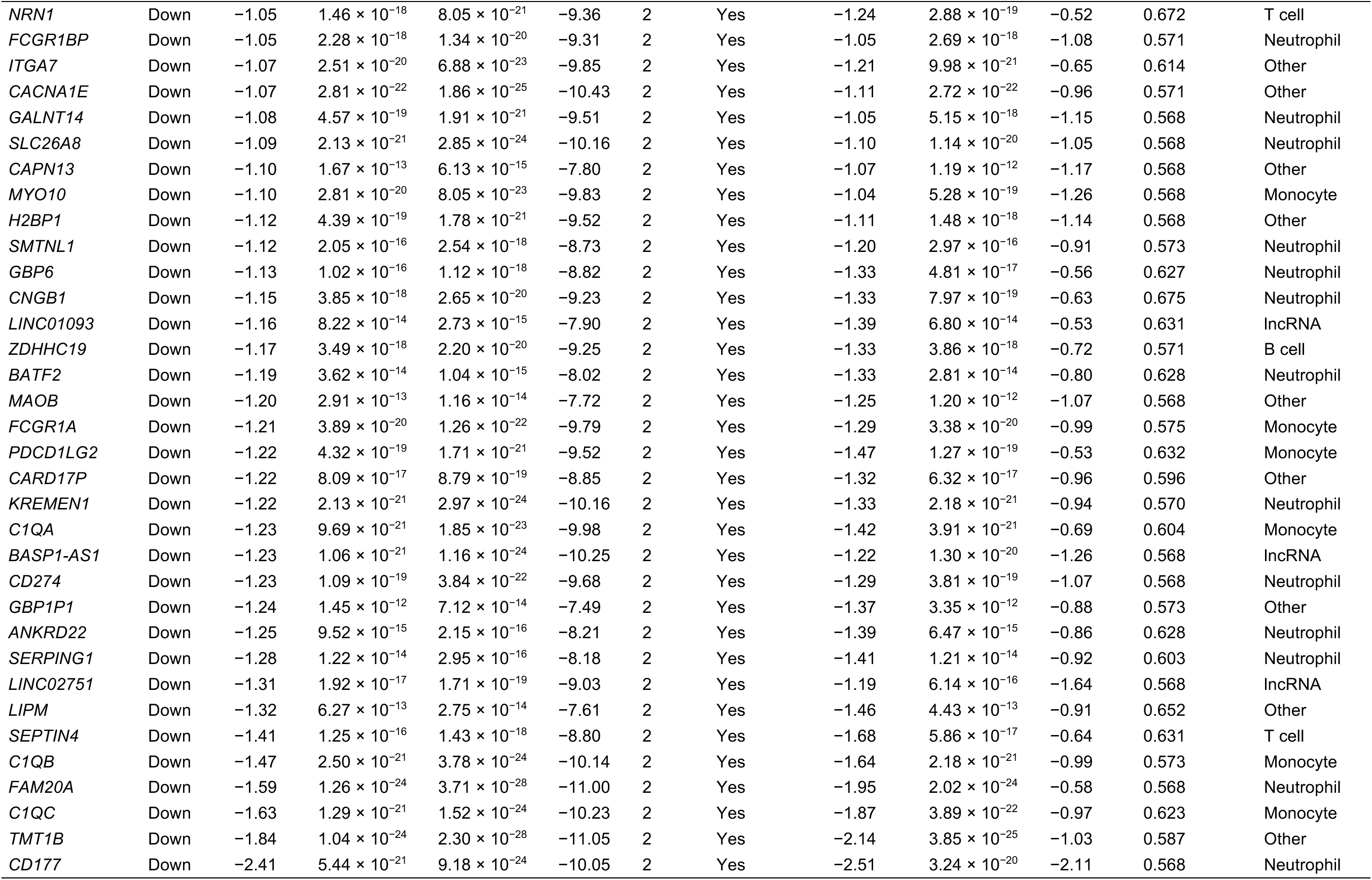
Meta-Analysis-derived DEGs with associated statistics. Gene symbol, direction of change, meta-analysis log2FC, Benjamini-Hochberg adjusted p-value, raw combined p-value, combined Stouffer Z-score, number of contributing studies, directional consistency flag, study-specific log2FC and adjusted p-value for GSE89403 and the study cohort, and Human Protein Atlas immune cell type assignment. Meta-DEGs are defined by FDR < 0.05, |log2FC| ≥ 1, and directional concordance across both cohorts.

| Gene symbol | Direction | Meta log<sub>2</sub>FC | Meta FDR | Meta p | Stouffer Z | Studies | Concordant | GSE89403 log<sub>2</sub>FC | GSE89403 FDR | Study cohort log<sub>2</sub>FC | Study cohort FDR | HPA cell type |
| --- | --- | --- | --- | --- | --- | --- | --- | --- | --- | --- | --- | --- |
| *ATP6V0CP4* | Up | 1.46 | $1.31 \times 10^{-15}$ | $2.06 \times 10^{-17}$ | 8.49 | 2 | Yes | 1.62 | $1.18 \times 10^{-15}$ | 1.02 | 0.596 | Other |
| *PRSS33* | Up | 1.43 | $1.93 \times 10^{-19}$ | $7.36 \times 10^{-22}$ | 9.61 | 2 | Yes | 1.64 | $2.76 \times 10^{-19}$ | 0.84 | 0.568 | Eosinophil |
| *ALOX15* | Up | 1.33 | $1.79 \times 10^{-19}$ | $6.69 \times 10^{-22}$ | 9.62 | 2 | Yes | 1.64 | $2.15 \times 10^{-20}$ | 0.47 | 0.713 | Eosinophil |
| *LINC02073* | Up | 1.32 | $1.18 \times 10^{-15}$ | $1.79 \times 10^{-17}$ | 8.51 | 2 | Yes | 1.52 | $1.35 \times 10^{-15}$ | 0.73 | 0.581 | lncRNA |
| *HRK* | Up | 1.29 | $1.64 \times 10^{-16}$ | $1.96 \times 10^{-18}$ | 8.76 | 2 | Yes | 1.42 | $4.98 \times 10^{-16}$ | 0.90 | 0.568 | B cell |
| *MEG3* | Up | 1.26 | $2.03 \times 10^{-13}$ | $7.64 \times 10^{-15}$ | 7.77 | 2 | Yes | 1.32 | $1.38 \times 10^{-12}$ | 1.11 | 0.568 | lncRNA |
| *CACNG6* | Up | 1.22 | $4.91 \times 10^{-17}$ | $4.90 \times 10^{-19}$ | 8.91 | 2 | Yes | 1.30 | $2.22 \times 10^{-16}$ | 1.01 | 0.568 | Basophil |
| *PLVAP* | Up | 1.21 | $6.76 \times 10^{-14}$ | $2.15 \times 10^{-15}$ | 7.93 | 2 | Yes | 1.37 | $6.96 \times 10^{-14}$ | 0.76 | 0.608 | Dendritic cell |
| *SIGLEC8* | Up | 1.20 | $2.53 \times 10^{-18}$ | $1.50 \times 10^{-20}$ | 9.29 | 2 | Yes | 1.55 | $9.73 \times 10^{-20}$ | 0.21 | 0.843 | Eosinophil |
| *SFRP2* | Up | 1.17 | $9.43 \times 10^{-13}$ | $4.44 \times 10^{-14}$ | 7.55 | 2 | Yes | 1.24 | $2.19 \times 10^{-12}$ | 0.97 | 0.573 | Other |
| *PLAAT5* | Up | 1.15 | $5.77 \times 10^{-18}$ | $4.36 \times 10^{-20}$ | 9.18 | 2 | Yes | 1.42 | $4.84 \times 10^{-19}$ | 0.40 | 0.773 | Eosinophil |
| *CXCL8* | Up | 1.06 | $4.76 \times 10^{-10}$ | $4.86 \times 10^{-11}$ | 6.58 | 2 | Yes | 0.88 | $2.22 \times 10^{-9}$ | 1.58 | 0.568 | Neutrophil |
| *PMP22* | Up | 1.05 | $5.79 \times 10^{-15}$ | $1.21 \times 10^{-16}$ | 8.28 | 2 | Yes | 1.19 | $9.44 \times 10^{-15}$ | 0.66 | 0.573 | Eosinophil |
| *LINC02972* | Up | 1.05 | $3.02 \times 10^{-14}$ | $8.40 \times 10^{-16}$ | 8.05 | 2 | Yes | 1.03 | $1.46 \times 10^{-13}$ | 1.10 | 0.568 | lncRNA |
| *SPNS3* | Up | 1.02 | $2.65 \times 10^{-18}$ | $1.62 \times 10^{-20}$ | 9.29 | 2 | Yes | 1.11 | $1.32 \times 10^{-17}$ | 0.75 | 0.568 | Eosinophil |
| *IL5RA* | Up | 1.02 | $6.12 \times 10^{-21}$ | $1.12 \times 10^{-23}$ | 10.03 | 2 | Yes | 1.32 | $3.54 \times 10^{-22}$ | 0.19 | 0.789 | Eosinophil |
| *MGAM2* | Down | -1.00 | $1.54 \times 10^{-11}$ | $1.01 \times 10^{-12}$ | -7.13 | 2 | Yes | -0.98 | $5.78 \times 10^{-11}$ | -1.09 | 0.568 | Other |
| *IGLV7-46* | Down | -1.01 | $8.23 \times 10^{-12}$ | $5.03 \times 10^{-13}$ | -7.22 | 2 | Yes | -0.90 | $1.71 \times 10^{-10}$ | -1.30 | 0.568 | B cell |
| *LINC02967* | Down | -1.01 | $1.39 \times 10^{-13}$ | $4.98 \times 10^{-15}$ | -7.83 | 2 | Yes | -1.12 | $1.08 \times 10^{-13}$ | -0.70 | 0.635 | lncRNA |
| *AIM2* | Down | -1.01 | $1.06 \times 10^{-21}$ | $1.13 \times 10^{-24}$ | -10.25 | 2 | Yes | -1.04 | $3.93 \times 10^{-21}$ | -0.93 | 0.568 | B cell |
| *SOCS3* | Down | -1.02 | $2.50 \times 10^{-21}$ | $3.86 \times 10^{-24}$ | -10.14 | 2 | Yes | -1.22 | $4.47 \times 10^{-22}$ | -0.45 | 0.667 | Neutrophil |
| *MCEMP1* | Down | -1.02 | $7.81 \times 10^{-19}$ | $3.90 \times 10^{-21}$ | -9.44 | 2 | Yes | -1.17 | $1.28 \times 10^{-19}$ | -0.59 | 0.684 | Monocyte |
| *ANXA3* | Down | -1.03 | $5.76 \times 10^{-16}$ | $8.04 \times 10^{-18}$ | -8.60 | 2 | Yes | -1.09 | $1.12 \times 10^{-15}$ | -0.87 | 0.570 | Neutrophil |
| *TLR5* | Down | -1.03 | $1.68 \times 10^{-25}$ | $1.54 \times 10^{-29}$ | -11.29 | 2 | Yes | -1.05 | $3.85 \times 10^{-25}$ | -0.99 | 0.568 | Monocyte |
| *C2* | Down | -1.03 | $9.16 \times 10^{-15}$ | $2.06 \times 10^{-16}$ | -8.22 | 2 | Yes | -1.14 | $6.57 \times 10^{-15}$ | -0.75 | 0.622 | Monocyte |
| *SHOC1* | Down | -1.03 | $2.44 \times 10^{-17}$ | $2.24 \times 10^{-19}$ | -9.00 | 2 | Yes | -0.92 | $4.35 \times 10^{-16}$ | -1.35 | 0.568 | Neutrophil |
| *CASP5* | Down | -1.04 | $8.59 \times 10^{-19}$ | $4.41 \times 10^{-21}$ | -9.42 | 2 | Yes | -1.16 | $4.84 \times 10^{-19}$ | -0.68 | 0.600 | Monocyte |
| *LINC03078* | Down | -1.05 | $1.67 \times 10^{-14}$ | $4.20 \times 10^{-16}$ | -8.13 | 2 | Yes | -1.01 | $2.31 \times 10^{-14}$ | -1.17 | 0.585 | lncRNA |
| *FGF13* | Down | -1.05 | $4.87 \times 10^{-19}$ | $2.11 \times 10^{-21}$ | -9.50 | 2 | Yes | -1.22 | $1.92 \times 10^{-19}$ | -0.59 | 0.614 | Neutrophil |

| Gene symbol | Direction | Meta<br>log<sub>2</sub>FC | Meta FDR | Meta p | Stouffer<br>Z | Studies | Concordant | GSE89403<br>log<sub>2</sub>FC | GSE89403<br>FDR | Study<br>cohort<br>log<sub>2</sub>FC | Study cohort<br>FDR | HPA cell<br>type |
| --- | --- | --- | --- | --- | --- | --- | --- | --- | --- | --- | --- | --- |
| *NRN1* | Down | -1.05 | $1.46 \times 10^{-18}$ | $8.05 \times 10^{-21}$ | -9.36 | 2 | Yes | -1.24 | $2.88 \times 10^{-19}$ | -0.52 | 0.672 | T cell |
| *FCGR1BP* | Down | -1.05 | $2.28 \times 10^{-18}$ | $1.34 \times 10^{-20}$ | -9.31 | 2 | Yes | -1.05 | $2.69 \times 10^{-18}$ | -1.08 | 0.571 | Neutrophil |
| *ITGA7* | Down | -1.07 | $2.51 \times 10^{-20}$ | $6.88 \times 10^{-23}$ | -9.85 | 2 | Yes | -1.21 | $9.98 \times 10^{-21}$ | -0.65 | 0.614 | Other |
| *CACNA1E* | Down | -1.07 | $2.81 \times 10^{-22}$ | $1.86 \times 10^{-25}$ | -10.43 | 2 | Yes | -1.11 | $2.72 \times 10^{-22}$ | -0.96 | 0.571 | Other |
| *GALNT14* | Down | -1.08 | $4.57 \times 10^{-19}$ | $1.91 \times 10^{-21}$ | -9.51 | 2 | Yes | -1.05 | $5.15 \times 10^{-18}$ | -1.15 | 0.568 | Neutrophil |
| *SLC26A8* | Down | -1.09 | $2.13 \times 10^{-21}$ | $2.85 \times 10^{-24}$ | -10.16 | 2 | Yes | -1.10 | $1.14 \times 10^{-20}$ | -1.05 | 0.568 | Neutrophil |
| *CAPN13* | Down | -1.10 | $1.67 \times 10^{-13}$ | $6.13 \times 10^{-15}$ | -7.80 | 2 | Yes | -1.07 | $1.19 \times 10^{-12}$ | -1.17 | 0.568 | Other |
| *MYO10* | Down | -1.10 | $2.81 \times 10^{-20}$ | $8.05 \times 10^{-23}$ | -9.83 | 2 | Yes | -1.04 | $5.28 \times 10^{-19}$ | -1.26 | 0.568 | Monocyte |
| *H2BP1* | Down | -1.12 | $4.39 \times 10^{-19}$ | $1.78 \times 10^{-21}$ | -9.52 | 2 | Yes | -1.11 | $1.48 \times 10^{-18}$ | -1.14 | 0.568 | Other |
| *SMTNL1* | Down | -1.12 | $2.05 \times 10^{-16}$ | $2.54 \times 10^{-18}$ | -8.73 | 2 | Yes | -1.20 | $2.97 \times 10^{-16}$ | -0.91 | 0.573 | Neutrophil |
| *GBP6* | Down | -1.13 | $1.02 \times 10^{-16}$ | $1.12 \times 10^{-18}$ | -8.82 | 2 | Yes | -1.33 | $4.81 \times 10^{-17}$ | -0.56 | 0.627 | Neutrophil |
| *CNGB1* | Down | -1.15 | $3.85 \times 10^{-18}$ | $2.65 \times 10^{-20}$ | -9.23 | 2 | Yes | -1.33 | $7.97 \times 10^{-19}$ | -0.63 | 0.675 | Neutrophil |
| *LINC01093* | Down | -1.16 | $8.22 \times 10^{-14}$ | $2.73 \times 10^{-15}$ | -7.90 | 2 | Yes | -1.39 | $6.80 \times 10^{-14}$ | -0.53 | 0.631 | lncRNA |
| *ZDHHC19* | Down | -1.17 | $3.49 \times 10^{-18}$ | $2.20 \times 10^{-20}$ | -9.25 | 2 | Yes | -1.33 | $3.86 \times 10^{-18}$ | -0.72 | 0.571 | B cell |
| *BATF2* | Down | -1.19 | $3.62 \times 10^{-14}$ | $1.04 \times 10^{-15}$ | -8.02 | 2 | Yes | -1.33 | $2.81 \times 10^{-14}$ | -0.80 | 0.628 | Neutrophil |
| *MAOB* | Down | -1.20 | $2.91 \times 10^{-13}$ | $1.16 \times 10^{-14}$ | -7.72 | 2 | Yes | -1.25 | $1.20 \times 10^{-12}$ | -1.07 | 0.568 | Other |
| *FCGR1A* | Down | -1.21 | $3.89 \times 10^{-20}$ | $1.26 \times 10^{-22}$ | -9.79 | 2 | Yes | -1.29 | $3.38 \times 10^{-20}$ | -0.99 | 0.575 | Monocyte |
| *PDCD1LG2* | Down | -1.22 | $4.32 \times 10^{-19}$ | $1.71 \times 10^{-21}$ | -9.52 | 2 | Yes | -1.47 | $1.27 \times 10^{-19}$ | -0.53 | 0.632 | Monocyte |
| *CARD17P* | Down | -1.22 | $8.09 \times 10^{-17}$ | $8.79 \times 10^{-19}$ | -8.85 | 2 | Yes | -1.32 | $6.32 \times 10^{-17}$ | -0.96 | 0.596 | Other |
| *KREMEN1* | Down | -1.22 | $2.13 \times 10^{-21}$ | $2.97 \times 10^{-24}$ | -10.16 | 2 | Yes | -1.33 | $2.18 \times 10^{-21}$ | -0.94 | 0.570 | Neutrophil |
| *C1QA* | Down | -1.23 | $9.69 \times 10^{-21}$ | $1.85 \times 10^{-23}$ | -9.98 | 2 | Yes | -1.42 | $3.91 \times 10^{-21}$ | -0.69 | 0.604 | Monocyte |
| *BASP1-AS1* | Down | -1.23 | $1.06 \times 10^{-21}$ | $1.16 \times 10^{-24}$ | -10.25 | 2 | Yes | -1.22 | $1.30 \times 10^{-20}$ | -1.26 | 0.568 | lncRNA |
| *CD274* | Down | -1.23 | $1.09 \times 10^{-19}$ | $3.84 \times 10^{-22}$ | -9.68 | 2 | Yes | -1.29 | $3.81 \times 10^{-19}$ | -1.07 | 0.568 | Neutrophil |
| *GBP1P1* | Down | -1.24 | $1.45 \times 10^{-12}$ | $7.12 \times 10^{-14}$ | -7.49 | 2 | Yes | -1.37 | $3.35 \times 10^{-12}$ | -0.88 | 0.573 | Other |
| *ANKRD22* | Down | -1.25 | $9.52 \times 10^{-15}$ | $2.15 \times 10^{-16}$ | -8.21 | 2 | Yes | -1.39 | $6.47 \times 10^{-15}$ | -0.86 | 0.628 | Neutrophil |
| *SERPING1* | Down | -1.28 | $1.22 \times 10^{-14}$ | $2.95 \times 10^{-16}$ | -8.18 | 2 | Yes | -1.41 | $1.21 \times 10^{-14}$ | -0.92 | 0.603 | Neutrophil |
| *LINC02751* | Down | -1.31 | $1.92 \times 10^{-17}$ | $1.71 \times 10^{-19}$ | -9.03 | 2 | Yes | -1.19 | $6.14 \times 10^{-16}$ | -1.64 | 0.568 | lncRNA |
| *LIPM* | Down | -1.32 | $6.27 \times 10^{-13}$ | $2.75 \times 10^{-14}$ | -7.61 | 2 | Yes | -1.46 | $4.43 \times 10^{-13}$ | -0.91 | 0.652 | Other |
| *SEPTIN4* | Down | -1.41 | $1.25 \times 10^{-16}$ | $1.43 \times 10^{-18}$ | -8.80 | 2 | Yes | -1.68 | $5.86 \times 10^{-17}$ | -0.64 | 0.631 | T cell |
| *C1QB* | Down | -1.47 | $2.50 \times 10^{-21}$ | $3.78 \times 10^{-24}$ | -10.14 | 2 | Yes | -1.64 | $2.18 \times 10^{-21}$ | -0.99 | 0.573 | Monocyte |
| *FAM20A* | Down | -1.59 | $1.26 \times 10^{-24}$ | $3.71 \times 10^{-28}$ | -11.00 | 2 | Yes | -1.95 | $2.02 \times 10^{-24}$ | -0.58 | 0.568 | Neutrophil |
| *C1QC* | Down | -1.63 | $1.29 \times 10^{-21}$ | $1.52 \times 10^{-24}$ | -10.23 | 2 | Yes | -1.87 | $3.89 \times 10^{-22}$ | -0.97 | 0.623 | Monocyte |
| *TMT1B* | Down | -1.84 | $1.04 \times 10^{-24}$ | $2.30 \times 10^{-28}$ | -11.05 | 2 | Yes | -2.14 | $3.85 \times 10^{-25}$ | -1.03 | 0.587 | Other |
| *CD177* | Down | -2.41 | $5.44 \times 10^{-21}$ | $9.18 \times 10^{-24}$ | -10.05 | 2 | Yes | -2.51 | $3.24 \times 10^{-20}$ | -2.11 | 0.568 | Neutrophil |

**Supplementary Table S2.** Differentially Expressed miRNAs with associated statistics. All 7 DE-miRNAs with base mean, log2FC (apeglm-shrunk), log2FC standard error, p-value, Benjamini-Hochberg adjusted p-value, and significance classification. Differential expression performed using DESeq2 with paired design (∼subject + condition) on study cohort small RNA-seq data (n = 11 pairs).

| miRNA | Base mean | log<sub>2</sub>FC | SE | p | FDR | Direction |
| --- | --- | --- | --- | --- | --- | --- |
| miR-151a-5p/151b | 9635.4 | -0.64 | 0.14 | $1.60 \times 10^{-7}$ | $8.33 \times 10^{-5}$ | Down |
| miR-139-5p | 849.7 | -0.75 | 0.23 | $5.20 \times 10^{-5}$ | 0.014 | Down |
| miR-144-5p | 142.4 | 0.94 | 0.36 | $2.13 \times 10^{-4}$ | 0.031 | Up |
| miR-28-3p | 719.6 | -0.81 | 0.32 | $3.24 \times 10^{-4}$ | 0.034 | Down |
| miR-144-3p | 452.2 | 1.09 | 0.56 | $4.27 \times 10^{-4}$ | 0.037 | Up |
| miR-32-5p | 56.2 | 1.07 | 0.52 | $5.09 \times 10^{-4}$ | 0.038 | Up |
| miR-4745-3p | 73.3 | -0.80 | 0.35 | $6.51 \times 10^{-4}$ | 0.042 | Down |

**Supplementary Table S3.** miRNA-mRNA target pairs identified by multiMiR. Each row represents a miRNA-mRNA pair supported by either validated experimental evidence or computational prediction in the multiMiR database. miRNA log₂FC and mRNA log₂FC denote log2 fold change (week 4 vs baseline) for the miRNA (DESeq2, study cohort) and mRNA (limma, study cohort), respectively. mRNA log₂FC (meta-analysis) is the meta-analysis log₂FC. Inverse: Yes indicates opposing directionality of miRNA and mRNA fold change, consistent with canonical miRNA-mediated repression.

| miRNA | Target | Evidence | multiMiR database(s) | miRNA log<sub>2</sub>FC | mRNA log<sub>2</sub>FC (study cohort) | mRNA log<sub>2</sub>FC (meta-analysis) | Inverse |
| --- | --- | --- | --- | --- | --- | --- | --- |
| miR-139-5p | *C1QB* | Validated | tarbase | -0.746 | -0.989 | -1.468 | No |
| miR-139-5p | *C2* | Validated | tarbase | -0.746 | -0.753 | -1.034 | No |
| miR-139-5p | *FGF13* | Validated | tarbase | -0.746 | -0.586 | -1.052 | No |
| miR-139-5p | *ITGA7* | Validated | tarbase | -0.746 | -0.654 | -1.067 | No |
| miR-139-5p | *KREMEN1* | Validated | tarbase | -0.746 | -0.936 | -1.223 | No |
| miR-139-5p | *MAOB* | Validated | tarbase | -0.746 | -1.074 | -1.201 | No |
| miR-139-5p | *MYO10* | Validated | tarbase | -0.746 | -1.261 | -1.102 | No |
| miR-139-5p | *PLAAT5* | Validated | tarbase | -0.746 | 0.400 | 1.155 | Yes |
| miR-139-5p | *PMP22* | Validated | elmmo; miranda; mirdb; mirtarbase; tarbase; targetscan | -0.746 | 0.657 | 1.049 | Yes |
| miR-139-5p | *SEPTIN4* | Validated | tarbase | -0.746 | -0.636 | -1.406 | No |
| miR-139-5p | *SERPING1* | Validated | tarbase | -0.746 | -0.919 | -1.279 | No |
| miR-144-5p | *HRK* | Validated | tarbase | 0.941 | 0.900 | 1.286 | No |
| miR-28-3p | *MYO10* | Validated | pita; tarbase | -0.813 | -1.261 | -1.102 | No |
| miR-32-5p | *CD274* | Validated | tarbase | 1.075 | -1.075 | -1.234 | Yes |
| miR-32-5p | *CXCL8* | Validated | tarbase | 1.075 | 1.580 | 1.061 | No |
| miR-32-5p | *FGF13* | Validated | tarbase | 1.075 | -0.586 | -1.052 | Yes |
| miR-32-5p | *MYO10* | Validated | pita; tarbase | 1.075 | -1.261 | -1.102 | Yes |
| miR-32-5p | *PMP22* | Validated | tarbase | 1.075 | 0.657 | 1.049 | No |
| miR-144-3p | *ANXA3* | Predicted | microcosm | 1.093 | -0.867 | -1.031 | Yes |
| miR-144-3p | *MAOB* | Predicted | diana_microt; miranda | 1.093 | -1.074 | -1.201 | Yes |
| miR-144-3p | *MGAM2* | Predicted | diana_microt | 1.093 | -1.086 | -1.004 | Yes |
| miR-144-5p | *FGF13* | Predicted | miranda | 0.941 | -0.586 | -1.052 | Yes |
| miR-28-3p | *C2* | Predicted | microcosm | -0.813 | -0.753 | -1.034 | No |
| miR-28-3p | *CD274* | Predicted | diana_microt | -0.813 | -1.075 | -1.234 | No |
| miR-4745-3p | *ITGA7* | Predicted | diana_microt | -0.802 | -0.654 | -1.067 | No |

**Supplementary Table S4.**
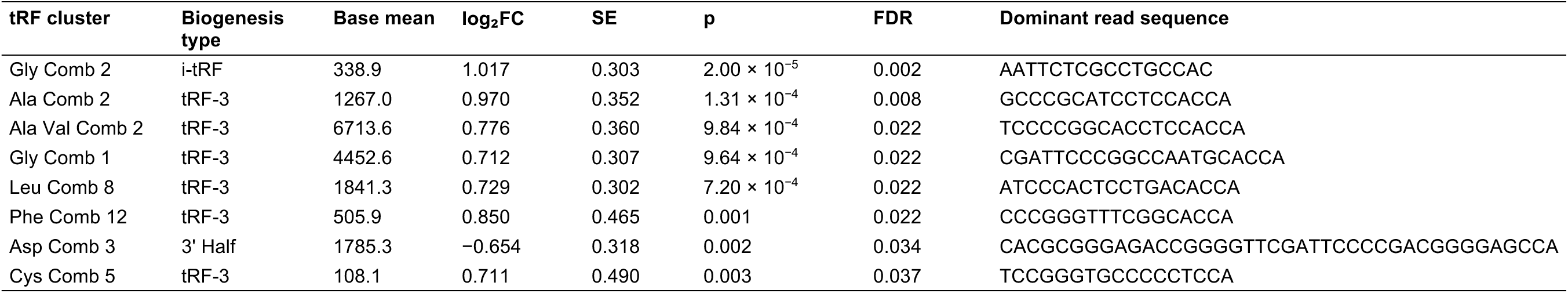
Differentially expressed tRFs with associated statistics and annotations. tRF cluster name, base mean, log2FC (apeglm-shrunk), log2FC standard error, p-value, Benjamini-Hochberg adjusted p-value, biogenesis type (tRF-3, i-tRF, or 3’ half) and dominant read sequence. All eight DE-tRFs were nuclear-encoded. Differential expression performed using DESeq2 with paired design on study cohort small RNA-seq data (n = 11 pairs).

**Supplementary Table S5.** Discrimination of week-4 sputum culture conversion by EARLY5 and published TB signatures. Baseline samples from GSE89403 (n = 76; 18 fast converters). EARLY5 scores are out-of-fold LOOCV predictions (LOOCV accuracy 0.724; sensitivity 0.44, specificity 0.81, using a majority of gene-pair votes). Sweeney3 and RISK6 were scored with ssGSEA using TBSignatureProfiler; higher scores in both were associated with slower conversion. AUC 95% CIs were calculated with the DeLong method. Paired DeLong tests compared each comparator with EARLY5 after aligning the direction of the scores.

**(a) Discrimination**
| Signature | n | AUC (95% CI) | Paired DeLong p vs EARLY5 |
| --- | --- | --- | --- |
| EARLY5 (LOOCV) | 76 | 0.761 (0.649–0.874) | — |
| Sweeney3 | 76 | 0.667 (0.520–0.814) | 0.171 |
| RISK6 | 76 | 0.628 (0.478–0.778) | 0.113 |

| n | True fast | True slow | TP | FN | FP | TN | Sensitivity | Specificity | PPV | NPV | Accuracy |
| --- | --- | --- | --- | --- | --- | --- | --- | --- | --- | --- | --- |
| 76 | 18 | 58 | 8 | 10 | 11 | 47 | 0.44 | 0.81 | 0.42 | 0.82 | 0.72 |

**Supplementary Table S6.** Cox proportional hazards models for time to sputum culture conversion. The event is sputum culture conversion, assessed at weeks 4, 8, 12 and 24; all participants converted by week 24. HR > 1 indicates faster conversion. Predicted fast/slow is the out-of-fold EARLY5 classification (fast n = 19, slow n = 57). Ties were handled with the Efron method. Proportional hazards assumptions were checked with Schoenfeld residuals (global p = 0.93, 0.32 and 0.21 for the unadjusted, adjusted and continuous-score models). With exact handling of ties, HRs were 2.71 (1.16–6.36; p = 0.022) unadjusted, 2.58 (0.90–7.39; p = 0.077) after adjustment for MGIT TTP, and 1.11 (0.83–1.49; p = 0.469) for the continuous score.

| Model | Term | n | HR (95% CI) | p | Ties |
| --- | --- | --- | --- | --- | --- |
| Unadjusted | Predicted fast vs slow | 76 | 1.85 (1.08–3.16) | 0.024 | Efron |
| Unadjusted (MGIT complete cases) | Predicted fast vs slow | 58 | 1.96 (1.06–3.65) | 0.033 | Efron |
| Adjusted for MGIT TTP | Predicted fast vs slow | 58 | 1.72 (0.90–3.31) | 0.102 | Efron |
| Adjusted for MGIT TTP | MGIT TTP (per IQR) | 58 | 1.30 (1.06–1.60) | 0.013 | Efron |
| Continuous score + MGIT TTP | EARLY5 score (per 0.2) | 58 | 1.04 (0.85–1.28) | 0.699 | Efron |
| Continuous score + MGIT TTP | MGIT TTP (per IQR) | 58 | 1.34 (1.05–1.70) | 0.017 | Efron |
| Unadjusted | Predicted fast vs slow | 76 | 2.71 (1.16–6.36) | 0.022 | Exact |
| Adjusted for MGIT TTP | Predicted fast vs slow | 58 | 2.58 (0.90–7.39) | 0.077 | Exact |
| Continuous score + MGIT TTP | EARLY5 score (per 0.2) | 58 | 1.11 (0.83–1.49) | 0.469 | Exact |

**Supplementary Table S7.** Stability of gene-pair selection across LOOCV folds. In each of 76 leave-one-out folds, k-TSP selected up to five gene pairs from the training data (k = 5 in 65 folds, k = 4 in 9, k = 3 in 2). The five pairs that make up EARLY5 were the most frequently selected.

| Gene pair | Folds selected (of 76) | % |
| --- | --- | --- |
| *DIS3L* > *DOCK4* | 76 | 100 |
| *COL9A2* > *FAM241A* | 72 | 95 |
| *PIEZO1* > *H2BC12* | 69 | 91 |
| *PDE4DIP* > *GRB10* | 67 | 88 |
| *PID1* > *LYPLA1* | 54 | 71 |
| *PID1* > *FCGR1BP* | 5 | 7 |
| *CYTH3* > *GRB10* | 4 | 5 |
| *POU2F2* > *CXCL16* | 3 | 4 |
| *RGS14* > *SIRPB1* | 3 | 4 |
| *XPC* > *RNF11* | 3 | 4 |
| *COL9A2* > *GCLM* | 2 | 3 |
| *ARHGEF2* > *SIRPB1* | 1 | 1 |
| *GGA2* > *H2BC12* | 1 | 1 |
| *KRT23* > *LYPLA1* | 1 | 1 |
| *PDE4DIP* > *DCUN1D3* | 1 | 1 |
| *PID1* > *ACSL3* | 1 | 1 |
| *PID1* > *NINJ2* | 1 | 1 |
| *POU2F2* > *SNX3* | 1 | 1 |
| *RTN1* > *ACSL3* | 1 | 1 |
| *XPO5* > *ACSL3* | 1 | 1 |

**Supplementary Table S8.** Immune cell type abundance at baseline and week 4 estimated by CIBERSORTx and MCP-counter. Mean estimated abundance at baseline and week 4 in GSE89403 (n = 86 pairs). CIBERSORTx values are LM22 cell fractions (samples with CIBERSORTx-p < 0.05); MCP-counter values are scores estimated from TPM. Changes were assessed with paired Wilcoxon signed-rank tests, with Benjamini-Hochberg FDR correction within each method. — indicates cell types estimated at zero in all samples, which were not tested.

| Method | Cell type | Mean baseline | Mean week 4 | Paired Wilcoxon p | FDR |
| --- | --- | --- | --- | --- | --- |
| CIBERSORTx | B cells naive | 0.0274 | 0.0205 | 0.001 | 0.005 |
| CIBERSORTx | B cells memory | 0.0314 | 0.0375 | 0.004 | 0.009 |
| CIBERSORTx | Plasma cells | 0.0039 | 0.0023 | $6.83 \times 10^{-5}$ | $3.41 \times 10^{-4}$ |
| CIBERSORTx | T cells CD8 | 0.0907 | 0.1381 | $4.45 \times 10^{-11}$ | $8.91 \times 10^{-10}$ |
| CIBERSORTx | T cells CD4 naive | 0.0213 | 0.0222 | 0.911 | 0.959 |
| CIBERSORTx | T cells CD4 memory resting | 0.0014 | 0.0015 | 0.514 | 0.683 |
| CIBERSORTx | T cells CD4 memory activated | 0.0063 | 0.0025 | 0.001 | 0.005 |
| CIBERSORTx | T cells follicular helper | 0.0015 | 0.0019 | 0.556 | 0.683 |
| CIBERSORTx | T cells regulatory (Tregs) | 0.0829 | 0.1005 | $1.04 \times 10^{-6}$ | $6.97 \times 10^{-6}$ |
| CIBERSORTx | T cells gamma delta | 0.0000 | 0.0000 | — | — |
| CIBERSORTx | NK cells resting | 0.0652 | 0.0786 | 0.004 | 0.009 |
| CIBERSORTx | NK cells activated | 0.0152 | 0.0180 | 0.079 | 0.132 |
| CIBERSORTx | Monocytes | 0.0335 | 0.0320 | 0.629 | 0.698 |
| CIBERSORTx | Macrophages M0 | 0.0001 | 0.0000 | 1.000 | 1.000 |
| CIBERSORTx | Macrophages M1 | 0.0009 | 0.0005 | 0.015 | 0.030 |
| CIBERSORTx | Macrophages M2 | 0.0193 | 0.0229 | 0.078 | 0.132 |
| CIBERSORTx | Dendritic cells resting | 0.0023 | 0.0021 | 0.536 | 0.683 |
| CIBERSORTx | Dendritic cells activated | 0.0039 | 0.0030 | 0.301 | 0.464 |
| CIBERSORTx | Mast cells resting | 0.0311 | 0.0297 | 0.581 | 0.683 |
| CIBERSORTx | Mast cells activated | 0.0000 | 0.0000 | — | — |
| CIBERSORTx | Eosinophils | 0.0000 | 0.0019 | 0.004 | 0.009 |
| CIBERSORTx | Neutrophils | 0.5616 | 0.4843 | $1.10 \times 10^{-8}$ | $1.10 \times 10^{-7}$ |
| MCP-counter | B lineage | 509.5 | 365.8 | $2.35 \times 10^{-5}$ | $4.71 \times 10^{-5}$ |
| MCP-counter | CD8 T cells | 48.3 | 66.9 | $4.39 \times 10^{-12}$ | $1.76 \times 10^{-11}$ |
| MCP-counter | Cytotoxic lymphocytes | 161.9 | 217.3 | $5.37 \times 10^{-8}$ | $1.43 \times 10^{-7}$ |
| MCP-counter | Monocytic lineage | 21.1 | 20.3 | 0.195 | 0.195 |
| MCP-counter | Myeloid dendritic cells | 4.5 | 4.7 | 0.160 | 0.183 |
| MCP-counter | Neutrophils | 536.8 | 493.0 | 0.003 | 0.003 |
| MCP-counter | NK cells | 4.2 | 4.9 | 0.002 | 0.003 |
| MCP-counter | T cells | 33.7 | 47.3 | $8.25 \times 10^{-15}$ | $6.60 \times 10^{-14}$ |

## Notes

### Competing Interest Statement

The authors have declared no competing interest.

